# Brain-behaviour modes of covariation in healthy and clinically depressed young people

**DOI:** 10.1101/487421

**Authors:** Agoston Mihalik, Fabio S. Ferreira, Maria J. Rosa, Michael Moutoussis, Gabriel Ziegler, Joao M. Monteiro, Liana Portugal, Rick A. Adams, Rafael Romero-Garcia, Petra E. Vértes, Manfred G. Kitzbichler, František Váša, Matilde M. Vaghi, Edward T. Bullmore, Peter Fonagy, Ian M. Goodyer, Peter B. Jones, NSPN Consortium, Raymond Dolan, Janaina Mourao-Miranda

## Abstract

Understanding how variations in dimensions of psychometrics, IQ and demographics relate to changes in brain connectivity during the critical developmental period of adolescence and early adulthood is a major challenge. This has particular relevance for mental health disorders where a failure to understand these links might hinder the development of better diagnostic approaches and therapeutics. Here, we investigated this question in 306 adolescents and young adults (14-24y, 25 clinically depressed) using a multivariate statistical framework, based on canonical correlation analysis (CCA). By linking individual functional brain connectivity profiles to self-report questionnaires, IQ and demographic data we identified two distinct modes of covariation. The first mode mapped onto an externalization/internalization axis and showed a strong association with sex. The second mode mapped onto a well-being/distress axis independent of sex. Interestingly, both modes showed an association with age. Crucially, the changes in functional brain connectivity associated with changes in these phenotypes showed marked developmental effects. The findings point to a role for the default mode, frontoparietal and limbic networks in psychopathology and depression.

## Introduction

Adolescence and early adulthood are periods of high risk for onset of many psychiatric disorders^1,2^, with up to a fifth of 18 to 25 year olds seeking professional help for psychological distress^3^. Despite this there are, as yet, no biological measures that inform early diagnosis and treatment. Neuroimaging techniques, especially functional Magnetic Resonance Imaging (fMRI)^4^, enable researchers to relate biological measures, such as patterns of functional brain connectivity, to the continuum of healthy to pathological states^5^. Here, we applied these techniques to uniquely identify underlying dimensions of brain-behaviour variation during a key developmental period.

Multivariate statistical methods^6^, such as canonical correlation analysis (CCA)^7^, allow an investigation of the link between multiple sets of measures, such as brain imaging and behavioural data, collected from the same individuals. Recently, an emerging number of studies report links between individual patterns of functional brain connectivity and item-level measures of behaviour and mental symptoms^8–11^. To the best of our knowledge no study has investigated this relationship in adolescents and young people, including those with mental health problems.

In the current study, we exploited resting-state fMRI (rsfMRI) and extensive item-level self-report questionnaire data, IQ and demographic information (that we refer as ‘behavioural data’ for simplicity) to investigate relationships between individual patterns of functional brain connectivity and individual sets of psychometrics/IQ/demographics. We used a sample of 281 healthy and 25 clinically depressed subjects, comprising adolescents and young adults (14-24y, 165 females) from the NeuroScience in Psychiatry Network (NSPN) study^12^.

All 306 subjects completed self-report questionnaires assessing well-being, affective symptoms, anxiety, impulsivity, compulsivity, self-esteem, self-harm, personality characteristics, psychotic spectrum symptoms, substance use, relations with peers and family and experience of trauma. These item-level measures were supplemented with measures of subjects’ fluid and crystallized intelligence as well as additional demographic information (age, gender and poverty index) amounting to a total of 364 behavioural (i.e. psychometrics/IQ/demographics) measures for each subject.

We acquired anatomical MRI scans and approximately 11 minutes of rsfMRI from the 306 subjects, which was then preprocessed as described in Methods. From the rsfMRI data, we extracted averaged time-series from 348 brain regions using subcortical regions from Freesurfer^13^ and the multi-modal parcellation of the Human Connectome Project (HCP)^14^. We then estimated functional brain connectivity for each individual using full correlation between all pairs of regional time-series. This resulted in a single connectivity profile (348 x 348 regions, i.e. 60378 connections) for each subject.

We used canonical correlation analysis (CCA)^7,8^ to investigate the relationships between brain connectivity and behaviour profiles across individuals. CCA is a multivariate analysis technique, which seeks maximal correlations between linear combinations of two or more sources of data (e.g. brain connectivity and behavioural data). To reduce the high dimensionality of both brain and behavioural data (60378 and 364 variables, respectively) we applied principal component analysis (PCA). We then performed a CCA resulting in pairs of canonical variates, which define modes of covariation between linear combinations of brain connectivity and behaviour profiles (in short ‘CCA modes’). On these canonical variates, subjects are represented by brain and behaviour scores, which describe the subject specific loadings in each CCA mode. We used a permutation approach^15^ to estimate both the optimal number of PCA components and the statistical significance of the CCA modes. Finally, we applied CCA embedded in a multiple hold-out framework^16^ to investigate the generalizability of the model, i.e. to assess whether the CCA mode represent associations that can be found on new data.

Given the age range of our sample, we expected a strong age (or developmental) effect on the brain-behaviour modes of covariation. We predicted that variation in these modes would also be related to the presence of depression^17^, given that our sample also included clinically depressed subjects. Finally, we hypothesised that psychopathological symptoms might be associated with a core set of abnormal functional brain networks incorporating default mode, frontoparietal and limbic networks as suggested by recent literature^17,18^.

## Results

We found two significant modes of covariation (Fig. 1) between patterns of functional brain connectivity and sets of behavioural measures, using a permutation approach (Supplementary Methods and Fig. S1). These were based on an optimal number of PCA components, *d* = 25, which explained 53% and 56% of the behaviour and brain connectivity variance, respectively. The first and second modes yielded canonical correlations of *q* = 0.62,*P*_*FWE*_ < 0.0001 and *q* = 0.58, *P*_*FWE*_ < 0.0134, respectively.

**Figure 1:**
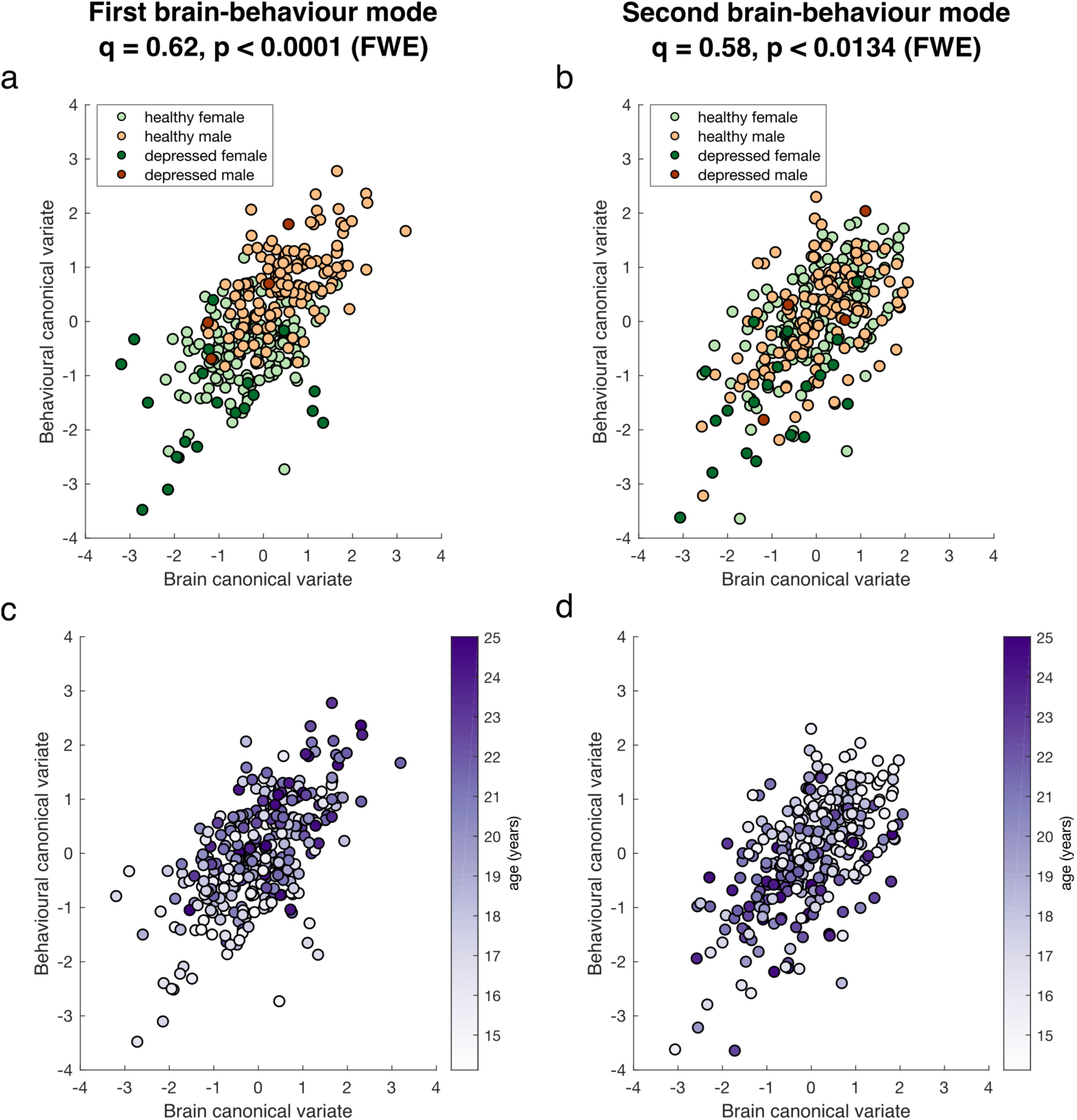
Significant brain-behaviour modes of covariation. Scatter plots showing the brain and behaviour scores for the first (**a** and **c**) and second (**b** and **d**) mode, where each dot represents an individual subject. Subjects are colour coded by: sex and clinical diagnosis (**a** and **b**); age (**c** and **d**). The canonical correlation, *q*, and corresponding p-value are shown on the top of each plot.

Figure 1 shows the two significant brain-behaviour modes of covariation, representing the correlation between brain and behaviour scores of individual subjects. The first mode is associated with sex and has an interaction with depression, with healthy males clustering towards higher scores and depressed females clustering towards lower scores (Fig. 1a). Additionally, younger adolescents can be seen to have lower scores whereas older ones are distributed more towards higher scores (Fig. 1c). The characteristics of the second mode were qualitatively different. Although depressed females seem to cluster towards lower scores (Fig. 1b) again, both males and females were evenly distributed along this mode, and younger adolescents having higher scores whereas older ones were more distributed towards lower scores (Fig. 1d).

To inform the association captured by each mode, we correlated the original behavioural and connectivity variables with the subjects’ brain and behaviour scores, respectively, and revealed the behavioural variables or brain connections most strongly associated with each CCA mode (Figs. 2-4). Figure 2a shows that the first CCA mode is positively associated with being male, age, measures of impulsivity, sensation seeking, drinking habits, and negatively associated with being female, depression-related symptoms and suicidal thoughts (for details, see Supplementary Table S1). Thus, the first mode has characteristic of an externalization/internalization axis, where extreme positive and negative scores represent vulnerability for males and females, respectively. Importantly, sex was weakly associated with the other top identified behavioural items (i.e. items most positively or negatively correlated with the behavioural variables) suggesting that those items are present due to an association with brain connectivity and not because of their association with sex (Supplementary Fig. S3). Brain connections most positively correlated with the first CCA mode (denoted by red edges in Fig. 3) included nodes within the dorsal and ventral attention networks and a somatomotor network; brain connections most negatively correlated (denoted by blue edges in Fig. 3) included nodes of the default mode, limbic and frontoparietal networks. Similar overall patterns were observed using different thresholds of top connections (Supplementary Fig. S4). In addition, when looking at the 0.5% most negatively correlated connections (top 302 connections), the subcortical network (mostly thalamus and caudate nucleus) also appeared negatively correlated with the first mode (including subcortical-subcortical connections and cortical connections with the default mode network, Supplementary Fig. S4). The list of 20 brain connections most positively/negatively associated with the first mode and their assignment to anatomical regions are described in Supplementary Table S2 and displayed on Supplementary Fig. S5.

**Figure 2:**
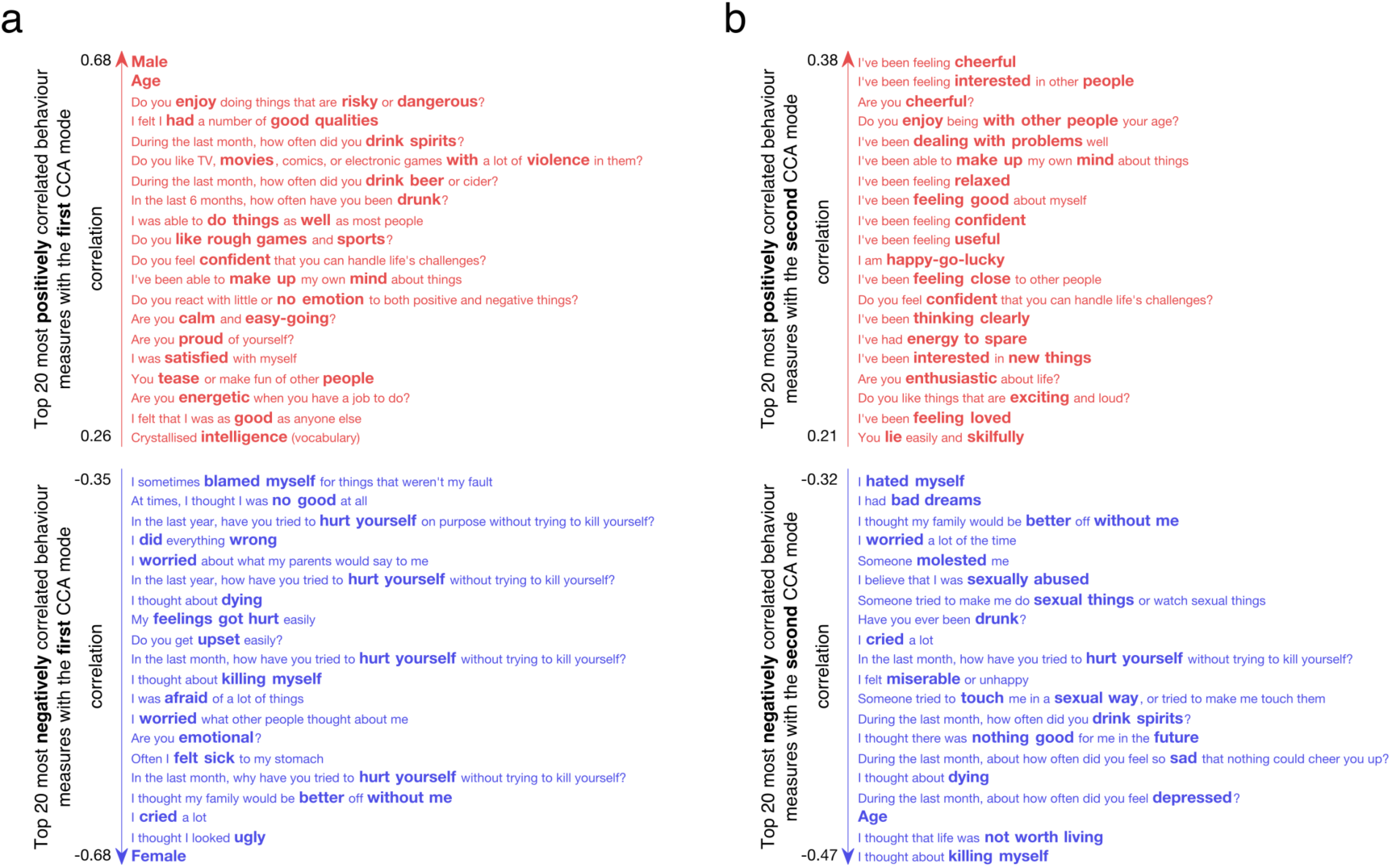
Correlations between the behavioural variables and the behavioural canonical variate (behaviour scores of all subjects) of the first (**a**) and second (**b**) CCA modes. Top 20 most positively and top 20 most negatively correlated variables are shown only. The exact correlation values and questionnaire items can be found in Supplementary Table S1 and Table S3.

**Figure 3:**
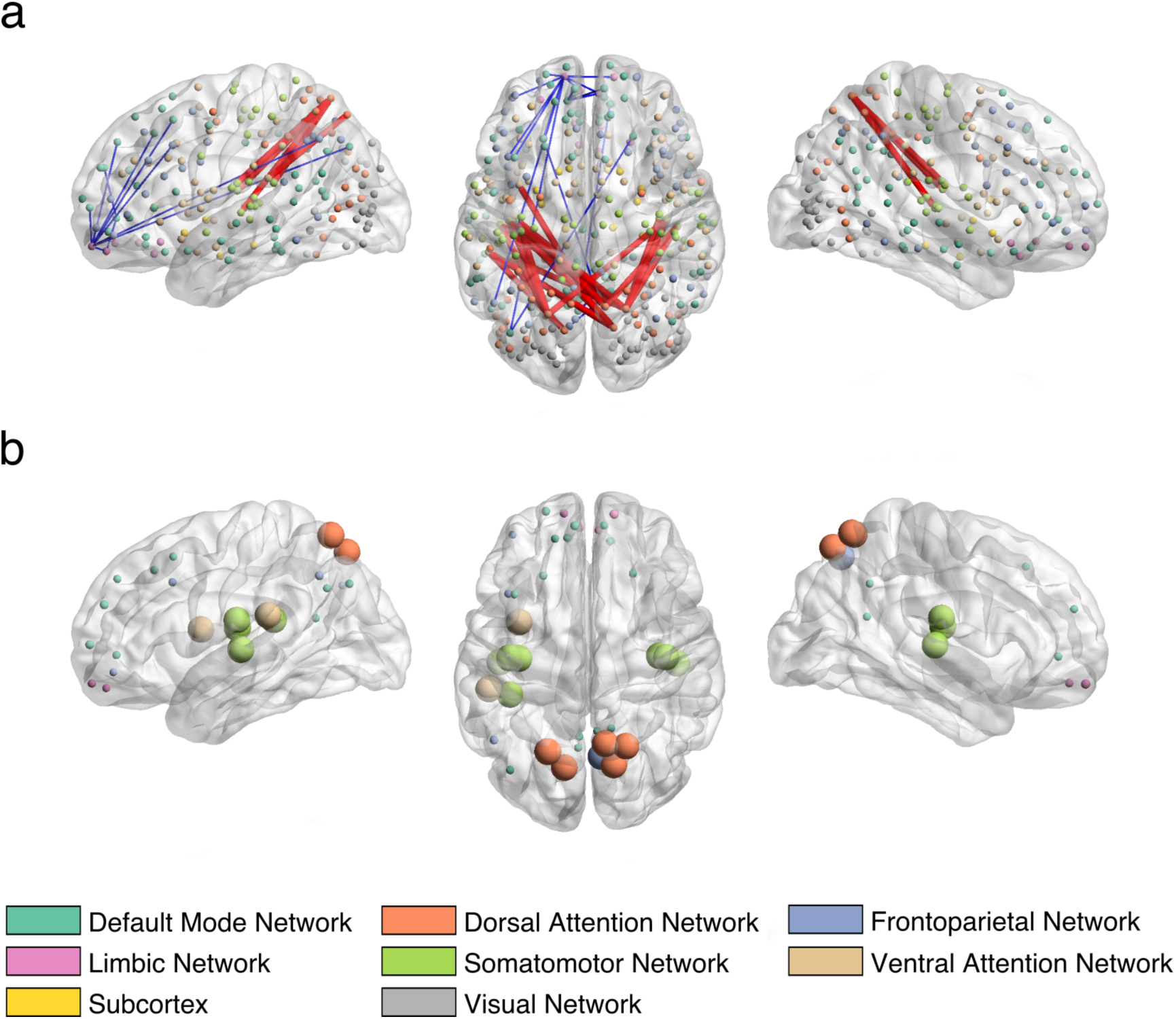
Correlations between the brain connectivity variables and the brain canonical variate (brain scores of all subjects) of the first CCA mode in sagittal (left and right) and axial views (middle). (**a**) Top 20 most positively and top 20 most negatively correlated brain connections. The thickness of the edges is proportional to the absolute correlation (red for positive correlations and blue for negative correlations). (**b**) Top 20 most positively and top 20 most negatively correlated brain connections summarized by nodes. Node size is proportional to the mean absolute correlation. Nodes are colour coded by resting state networks assigning each node to one of 7 cortical networks (based on the maximal surface based intersection) described in Yeo et al 2011^44^ or the subcortex. The full list of correlation values and respective labels can be found in Supplementary Table S2.

For the second mode, the most positively correlated behavioural variables (Fig. 2b) related to measures of mental well-being, self-esteem, confidence, while the most negatively associated related to age, depression-related symptoms, drinking habits, suicidal thoughts and sexual abuse. Thus, this second mode captures a well-being/distress axis, along which individuals vary from high mental well-being through to distress (for details, see Supplementary Table S3). The brain connections most positively correlated (depicted in red edges in Fig. 4) with this CCA mode included nodes involving mainly the default mode and subcortical networks (thalamus); brain connections most negatively correlated (depicted in blue edges in Fig. 4) included nodes within the dorsal and ventral attention networks and the visual and somatomotor networks. A largely similar overall pattern of networks was observed using different thresholds of top connections (Supplementary Fig. S4). In addition, when looking at the 0.5% most positively correlated connections (top 302 connections), the limbic and frontoparietal networks also appeared positively correlated with the second mode (including cortico-cortical connections and subcortical connections mostly with the thalamus, putamen and accumbens nucleus, Supplementary Fig. S4). The list of 20 brain connections most positively/negatively associated with the second mode and their assignment to anatomical regions are described in Supplementary Table S4 and displayed on Supplementary Fig. S6.

**Figure 4:**
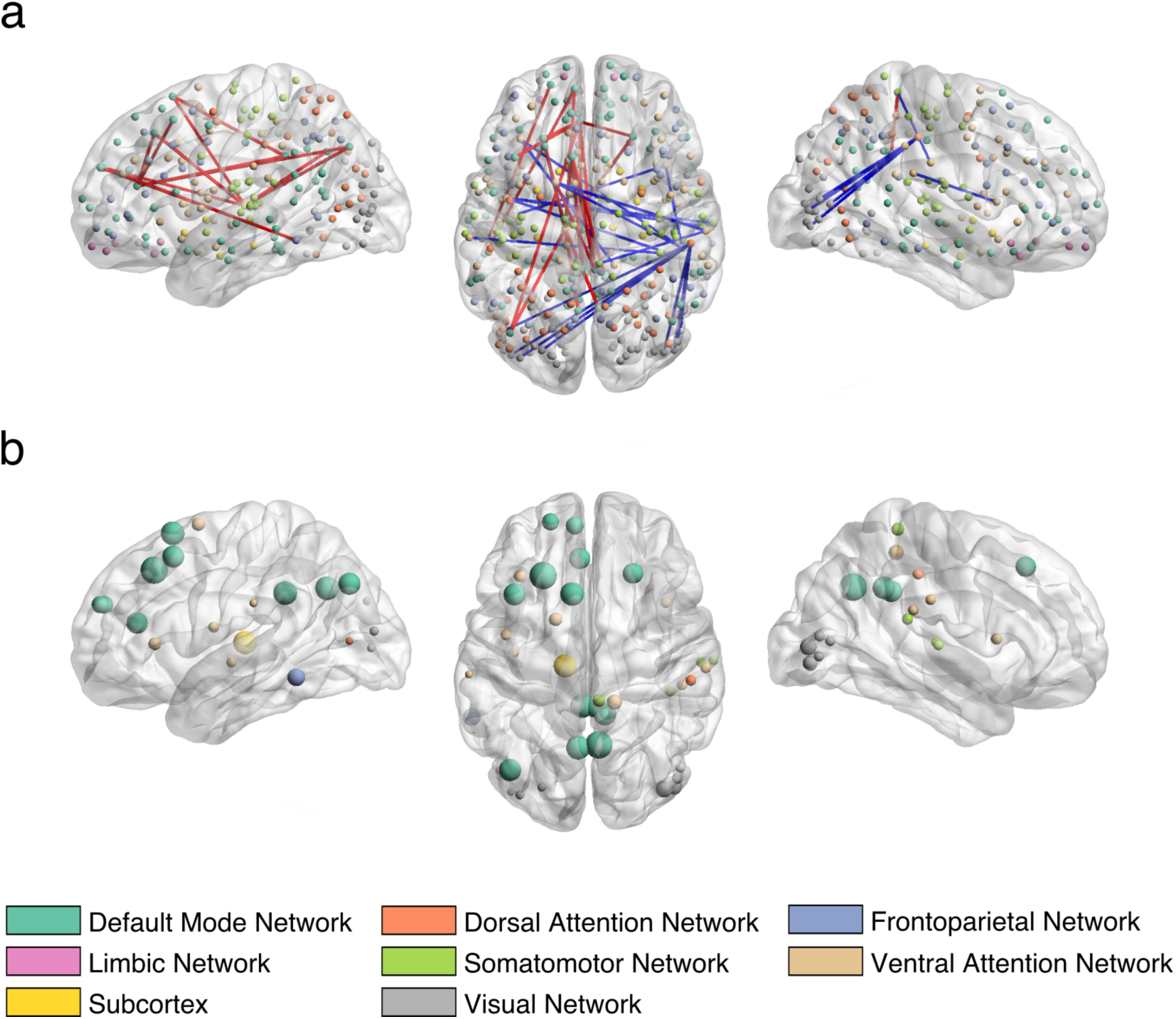
Correlations between the brain connectivity variables and the brain canonical variate (brain scores of all subjects) of the second CCA mode in sagittal (left and right) and axial views (middle). (**a**) Top 20 most positively and top 20 most negatively correlated brain connections. The thickness of the edges is proportional to the absolute correlation (red for positive correlations and blue for negative correlations). (**b**) Top 20 most positively and top 20 most negatively correlated brain connections summarized by nodes. Node size is proportional to the mean absolute correlation. Nodes are colour coded by resting state networks assigning each node to one of 7 cortical networks (based on the maximal surface based intersection) described in Yeo et al 2011^44^ or the subcortex. The full list of correlation values and respective labels can be found in Supplementary Table S4.

As validation of our model, we also applied a multiple hold-out framework (Supplementary Methods and Fig. S2) and found one brain-behaviour mode of covariation (Supplementary Fig. S7), based on ten PCA components (*d* = 10), which explained 40% and 47% of the behaviour and brain connectivity variance, respectively. Importantly, the distribution of subjects along the CCA main axis showed the same trend in the training and testing sets (Supplementary Fig. S8). When ranking the original behavioural and connectivity variables according to their correlation with the subjects’ brain and behaviour scores, we obtain a very similar overall ranking for both the permutation and hold-out frameworks (Supplementary Fig. S9). However, the overlap is not very large when we consider alone the top 20 most positively/negatively correlated behavioural and brain variables (Supplementary Tables S1-2 and Tables S5-6). Indeed, the scatter plots in Supplementary Fig. S9 show that only 11 brain connectivity and 11 behavioural variables overlapped when those top variables were chosen. This might be explained by the fact that we are just displaying the very top variables ranked according to their correlation value, and some correlation values simply differ from each other on the fourth decimal place. However, we can see that the overlap is more extended when the top 5% most positively/negatively correlated variables are selected (Supplementary Fig. S9).

## Discussion

In summary, leveraging both resting-state fMRI and behavioural data within a multivariate analysis framework, we identified two brain-behaviour modes of covariation in a sample of 306 adolescents and young adults. The first CCA mode relates to an externalization/internalization axis, which is associated with sex. Specifically, it suggests that males are more susceptible to disruptive behaviour and alcohol use, whilst females are more susceptible to depression and self-harm. The second CCA mode relates to a well-being/distress axis which covers positive symptoms of well-being on one side and negative symptoms related to depression, suicidal thoughts, history of sexual abuse and alcohol use on the other side. Both modes are also associated with age, which could be expected considering that the sample age range covers an important developmental period. Importantly, the brain networks related to both CCA modes align well with models of brain development highlighting the sequential maturation of subcortical and cortical regions in adolescence^19,20^ and models of psychopathology^17,18^.

Both CCA modes are conceptually associated with broadly described depressive psychopathology and can hence be seen as helping refine this clinical concept. It is therefore important to understand whether they capture distinctions in brain connectivity profiles alone or capture also distinctions in descriptive psychopathology. At first glance, the behavioural items common to both modes of depression, such as e.g. “…life was not worth living”, “I thought about dying”, “I cried a lot” might support the former hypothesis. Nevertheless, there are three clear differences:

Firstly, the first mode is associated with a more anxious, agitated and behaviourally-activated expression of depression (four self-harm items, “I felt sick…”, “I worried…”, “I was afraid…”, “Are you emotional?”). Conversely, the second mode is associated with a more anhedonic and amotivational state (positively correlated with “…life was not worth living”, “…nothing good for me in the future”, “…feel so sad…” and negatively correlated with “…feeling interested in other people”). Interestingly, similar ‘anxious’ and ‘anhedonic’ axes have been found in other large data-driven depression studies^9,21^.

Secondly, the first CCA mode is strongly correlated with sex, but the second mode is not. Thus the latter is a more sex independent dimension of psychopathology. Also, depression-related variables of the first mode are associated with younger age, whilst depression-related variables of the second mode are associated with older age (Fig. 1c-d and Fig. 2). Accordingly, depression in the first CCA mode is related to behavioural items, such as e.g. “…I looked ugly”, “…my family would be better off without me”, “I worried about what my parents would say…”, which are more likely to be hallmarks of depression at a younger age. On the contrary, distress in the second CCA mode is related to items thought to characterise depression at an older age (e.g. “I thought about killing myself”, being drunk and drinking spirits).

Thirdly, depression in the second mode is associated with sexual abuse and is negatively associated with feeling loved, confident and close to other people, perhaps indicating that sexual abuse affects these traits (although causal attributions are not possible in this dataset).

The strong relationship between sex and the first CCA mode is striking in light of recent findings that there is <10% overlap in gene expression changes in the brains of male and female humans with depression – at least in the prefrontal cortex and insula (other cortical areas were not sampled)^22^. Moreover, the authors demonstrated that a similar lack of overlap between the sexes also exists in a chronic variable stress mouse model^22^. It is interesting that both insula and the prefrontal cortex dominate the connections of the first CCA mode, being either positively (insula) or negatively (prefrontal cortex) correlated with depression. This suggests that sex interacts with depression risk in these (and likely other) areas in a way that might be fundamental to the disorder.

Adolescence and early adulthood is the peak age of onset for many psychiatric disorders^1,2^, rendering understanding vulnerability of individuals at this age is of particular relevance. Importantly, most items correlated with the CCA modes are related to psychopathology, and so the identified CCA modes might represent a two-dimensional space not only related to current depressive symptoms (or their absence), but to a latent vulnerability to psychopathology. Deeper understanding of this vulnerability may powerfully inform biologically informed interventions in young people^23^.

Substance use is highly correlated with psychiatric disorders^24,25^, and it is especially detrimental in adolescence. Personality traits have an etiological role in the development of alcohol and substance use, and a vast body of research implicates two broad personality domains with opposing action tendencies, namely inhibition and disinhibition^26,27^. Our results concur with such a model. Alcohol usage is associated with both of our CCA modes in opposing directions. Behavioural items resembling to a disinhibited personality (first CCA mode) are positive correlations with e.g. “…enjoy doing things that are risky and dangerous?”, “…like TV, movies, comics, or electronic games with a lot of violence in them?” or “Do you like rough games and sports?”; whereas items suggestive of an inhibited personality (second CCA mode) are negative correlations with e.g. being interested in or enjoying the company of other people, or being interested in new things.

As discussed above, age was associated with both CCA modes. The first CCA mode correlated positively with age (depicted in red in Fig. 2), attentional and frontoparietal networks (depicted in red in Supplementary Fig. S4) and negatively correlated with subcortical-subcortical connections as well as connections within the limbic system (depicted in blue in Supplementary Fig. S4). These results are consistent with models of adolescent brain development, demonstrating that subcortical and limbic regions mature in early adolescence followed by the maturation of cortico-cortical connections^19,20^. The second CCA mode was negatively correlated with age (depicted in blue in Fig. 2), connections within and between attentional networks (depicted in blue in Supplementary Fig. S4) and was positively correlated with various subcortical-cortical connections (depicted in blue in Supplementary Fig. S4). Again, these results corroborate the aforementioned models of adolescent brain development. In particular, the results of the two CCA modes substantiate the sequential maturation of brain circuits, namely, the fine-tuning of circuits from subcortical-subcortical (early adolescence) to cortico-subcortical (late adolescence) and cortico-cortical (young adulthood)^28^. Furthermore, the sequential maturation of brain circuits might be a risk factor for alcohol use^29^, which aligns well with the strong positive correlation between alcohol use and age found in both CCA modes (Fig. 2).

Our connectivity results are also consistent with recent literature suggesting that most psychiatric disorders emerge as a result of impairments within a few core brain circuits and networks^10,17,18^. In particular, the first mode was negatively correlated with depression and connections of the default mode, frontoparietal and limbic networks (Fig. 3); whilst the second mode was negatively correlated with depression and positively correlated with many default mode areas (Fig. 4). These networks underlie core social, executive and affective cognition, respectively, and dysfunctions in these networks might result in specific domains of symptoms (e.g. alterations in default mode network connectivity resulting in impaired self-representation and social functioning)^17^. Interestingly, due to the strong interplay between these networks, the aberrant functioning in any of these could cause impairments of the others. For example, excessive coupling between the limbic and default mode networks could mean that initial dysfunction in the former may propagate to the latter, causing depressive symptoms^10,30,31^. Conversely, a default mode network that can only dominate but cannot reciprocally communicate with the limbic network could prevent positive mood being established by the latter^32^.

It is common practice in statistics to estimate a model using the entire dataset, once model selection has been performed using the permutation-based approach as described above. In contrast, in machine learning a dataset is often split into train and test sets (or train/test/validation sets) using cross-validation procedures, such that the model parameters are estimated on a training data and the model performance is estimated on a testing data. The permutation-based approach may be preferable to cross-validation when the number of samples is not very large, since it avoids the need to split the data into even smaller sets for training and testing. However, cross-validation approaches might be preferable if one wants to measure the robustness and generalisability of the results. In the present study we also ran CCA embedded in a multiple hold-out framework (Supplementary Methods and Fig. S2) which was proposed by Monteiro et al^16^. We found one mode of covariation, which was comparable to the first one found using the permutation framework (Supplementary Fig. S7). The second mode obtained using the permutation framework was not found with the hold-out framework, potentially due to the small sample size and the strictness of the multiple hold-out framework. The most striking finding obtained with this approach was that the distribution of the subjects along the CCA main axis on the testing set was very similar to the training set (Supplementary Fig. S8), which means that the CCA mode generalised for the test set.

Finally, we acknowledge limitations to the current study. Methodological limitations relate to the pipeline choice, which includes use of an atlas, full correlation as a connectivity metric and the PCA dimensionality reduction step that might remove a significant amount of signal variability of potential interest. Further work exploring other approaches to estimating resting state connectivity (e.g. Independent Component Analysis (ICA)^8^ and partial correlation^33^) and regularized or sparse CCA could be investigated to overcome potential limitations of the current pipeline. We also acknowledge the cross-sectional nature of our analysis. Future studies involving longitudinal samples could investigate how the described brain-behaviour modes of covariation change over time or whether they are predictive of future psychopathology. In addition, although we have used a multiple hold-out framework, we should ideally use an independent replication sample to validate our model. Finally, there are limitations of sample size. Future studies including big datasets, such as the ABCD study^34^, will be useful to explore higher variability in general population and potentially find different dimensions of psychopathology or groups of at risk adolescents.

In conclusion, our results demonstrate that identifying brain-behaviour modes of covariation in healthy and depressed young people provides a powerful way of understanding the latent dimensions underlying abnormal mental states and behaviour^35^ and brings potential new insights into the mediation of vulnerability to mental health disorders.

## Supporting information

## Methods

### Subjects

In total, 2406 healthy subjects and 50 subjects clinically diagnosed with depression (diagnosis and referral made by the subject’s NHS care service) aged 14 to 24 years were recruited from schools, colleges, National Health Service (NHS) primary care and mental health services, and via direct advertisement in London and Cambridgeshire^12^. This was carried out by the University College London and University of Cambridge NeuroScience in Psychiatry Network (NSPN) research initiative, supported by a strategic award from the Wellcome Trust. A Magnetic Resonance Imaging (MRI) cohort was subsampled from the primary cohort, comprising a healthy cohort of 318 subjects and a depression cohort of 37 subjects. Furthermore, a demographically balanced cohort of 297 subjects was subsampled from the healthy cohort, with approximately 60 subjects in each of five age-defined strata: 14–15 years inclusive, 16–17, 18–19, 20–21, and 22–24 years.

Of the healthy cohort (n=297), 2 subjects were excluded due to low quality images, 1 was excluded due to gross radiological abnormalities, 4 were excluded due to missing convergence in ME-ICA pre-processing, and 9 were excluded due to excessive motion during the resting-state functional scan (5 subjects with maximum framewise displacement larger than 1.3 mm and 4 subjects with mean framewise displacement of 0.3 mm using calculation by Power et al 2012^36^). Of the depression cohort (n=37), 3 subjects were excluded due to low quality anatomical scans, 1 was excluded due to radiological artefacts, 4 were excluded due to motion-induced Freesurfer reconstruction errors, 1 was excluded due to lack of convergence in ME-ICA pre-processing, 1 was excluded due to extremely low explained variance in ME-ICA pre-processing (<20%) and 2 were excluded due to excessive motion during the resting-state functional scan (applying the same criteria as for the healthy cohort). These exclusion criteria produced a final healthy cohort consisting of 281 subjects (mean age=19.13, SD=2.88, 144 females) and a final depression cohort comprising 25 subjects (mean age=16.80, SD=1.15, 21 females).

Written, informed consent was obtained for all subjects over the age of 16 years. For subjects under the age of 16, written informed assent was obtained for the subject and written informed consent from their parent/legal guardian. The study was ethically approved by the Cambridge Central Research Ethics Committee and conducted in accordance with NHS research governance standards.

### Behavioural and demographic data

Subjects completed self-report questionnaires and cognitive tests as part of the NSPN data acquisition^12^. We used the following subset of these measures at the baseline study visit that assess psychopathological symptoms, personality characteristics, mental well-being and IQ as follows: Antisocial Behaviours Checklist (ABQ); Antisocial Process Screening Device (APSD); Barratt Impulsivity Scale (BIS); Child and Adolescent Dispositions Scale (CADS); Child Trauma Questionnaire (CTQ); Drugs Alcohol and Self-Injury (DASI); Inventory of Callous-Unemotional Traits (ICU); Kessler Psychological Distress Scale (K10); Leyton Obsessional Inventory (LOI); Moods and Feelings Questionnaire (MFQ); Revised Children’s Manifest Anxiety Scale (RCMAS); Rosenberg Self-Esteem Scale (SES); Schizotypal Personality Questionnaire (SPQ); Wechsler Abbreviated Scale of Intelligence (WASI); Warwick Edinburgh Mental Wellbeing Scale (WEMWBS).

Finally, we added four demographic variables (age, sex, and socioeconomic deprivation index; hot coding was used for sex resulting in 2 variables and see Supplementary Material for the calculation of the deprivation index) to the items of these questionnaires resulting in a total of 372 variables.

### MRI data acquisition

All MRI data were acquired on three identical 3T whole-body MRI systems (Magnetom TIM Trio; VB17 software version; Siemens Healthcare): two located in Cambridge and one located in London. Between-site reliability and tolerability of all MRI procedures were satisfactorily assessed by a pilot study of five healthy volunteers at each site^37^. Only scans at the baseline visit were included in the current study. Structural MRI scans were acquired using multi-echo acquisition protocol with six equidistant echo times (TE) between 2.2 and ms and averaged to form a single image of increased signal-to-noise ratio (SNR)^37^. Apparent longitudinal relaxation rate R1 (R1=1/T1w) was calculated using previously developed models to create quantitative R1 maps^38–40^. Other acquisition parameters included: temporal resolution (TR) of 18.70 ms, spatial resolution 1.0 mm isotropic, field of view (FOV) = 256 x 256, 176 sagittal slices and parallel imaging using GRAPPA factor 2 in anterior-posterior phase-encoding (PE) direction. Resting-state fMRI (rsfMRI) data were acquired using multi-echo acquisition protocol with three echo times TE = 13, 31, 48 ms, temporal resolution (TR) of 2420 ms, spatial resolution 3.8 mm isotropic with 10% gap, sequential slice acquisition, FOV = 240 x 240 mm, 34 oblique slices; bandwidth 1/4 2368 Hz/pixel and matrix size = 64 x 64 x 34.

### Structural MRI pre-processing

R1 images were used to perform a surface reconstruction of each subject using Freesurfer *recon-all*^41^ (https://surfer.nmr.mgh.harvard.edu/). Freesurfer average subject (fsaverage) was parcellated using a multimodal scheme that subdivides the cortex into 360 bilaterally symmetric regions based on Human Connectome Project (HCP) data^14^. HCP parcellation was transformed from fsaverage space to the cortical surface of each individual subject using Freesurfer *mri_surf2surf*. In addition, 16 regions were used from the subcortical segmentation of Freesurfer (thalamus-proper, caudate, putamen, pallidum, hippocampus, amygdala, accumbens-area, ventral diencephalon (DC) for each hemisphere).

### Resting-state fMRI pre-processing

rsfMRI data were pre-processed using multi-echo independent component analysis^42,43^ (ME-ICA). ME-ICA identifies BOLD components that scale linearly with TE and discards remaining components to reduce motion-related artefacts. Only BOLD components were optimally combined to generate the denoised time-series of each voxel. A wavelet filtering was used to focus on the physiologically relevant frequency range of 0.025-0.111 Hz (scales 2 and 3). Functional scans were coregistered with each individual structural R1 images for time-series extraction. Regional time-series were estimated as the average time-series of all the voxels included in each of the 360 cortical and 16 subcortical regions. 28 regions (mostly near the frontal and temporal pole) were excluded due to low regional mean signal in at least one subject (*z*-score across regions within subject, *Z* < –1.96), resulting in a total of 348 retained regions. Functional connectivity was calculated as the pairwise Pearson-correlation between time-series of each possible pair of regions resulting in a total of 60378 brain variables per subject.

### Behaviour and demographic data processing

The initial considered questionnaire data comprised 372 item level variables. However, we removed 8 variables for which more than 95% of subjects had the same value. Missing data were replaced by the median of the respective variable across subjects. This resulted in a total of 364 behaviour variables per subject.

### Confounds

We identified two main confound measures, whose effect was not of interest and were therefore regressed from both brain and behavioural data:

1. Mean framewise displacement: a summary statistic quantifying average subject head motion during the rsfMRI acquisitions (using calculation by Power et al 2012^36^).
2. Site: each MRI site (two in Cambridge and one in London) was encoded as a one-hot variable (for example: [0 0 1] for London).

Finally, each brain and behavioural variable was mean-centred and normalised (*μ* = 0, *σ* = 1), and these brain (*X* ∈ ℝ^*n*×*p*^, where *n* is the number of subjects and *p* is the number of brain connectivity variables) and behaviour (*Y* ∈ ℝ^*n*×*q*^, where *n* is the number of subjects and *q* is the number of behavioural variables) matrices were then entered into the CCA analysis.

### Canonical correlation analysis

We used Canonical Correlation Analysis (CCA) to investigate modes of covariation between patterns of brain connectivity and behavioural data. CCA seeks maximal correlations between linear combinations of two or more sources of data, e.g. brain connectivity and behavioural data.

To be able to apply CCA directly on this dataset, where the number of variables in both connectivity and behavioural data is higher than the number of subjects, we first reduced the dimensionality of the data using principal component analysis (PCA). In summary, the pre-processed brain and behavioural data matrices, *X* ∈ ℝ^*n*×*p*^and *Y* ∈ ℝ^*n*×*p*^, respectively, were first decomposed into *X*_*d*_ ∈ ℝ^*n*×*d*^ and *Y*_*d*_ ∈ ℝ^*n*×*d*^, where d is the number of PCA components. We chose the optimal number of components based on the permutation approach described below, using 9 different PCA dimensionalities (*d* = 5,10,25,50,75,100,125,150,200).

After the dimensionality reduction step, *X*_*d*_ and *Y*_*d*_ are fed into CCA, which outputs d components (modes of covariations, or in short, ‘CCA modes’). Each CCA mode includes a pair of canonical basis vectors *u* ∈ ℝ^*p*×*1*^ and *v* ∈ ℝ^*q*×*1*^ indicating the direction of maximum brain-behaviour correlation; as well as a pair of canonical variates *p*_*X*_ ∈ ℝ^*n*×*1*^ and *p*_*Y*_ ∈ ℝ^*n*×*1*^ obtained by projecting *X*_*d*_ and *Y*_*d*_ onto their corresponding canonical basis vectors. On these canonical variates, subjects are represented by brain and behaviour scores, which describe the subject specific loadings in each CCA mode. Significance for the CCA modes was assessed based on the permutation framework described below. To find brain connections and behavioural variables most strongly associated with the CCA modes, we correlated the pre-processed brain and behaviour variables with the canonical variates *p*_*X*_ and *p*_*Y*_, respectively. Then for each significant CCA mode, we have a vector of correlations for the top brain connectivity variables (Figs. 3-4) and a vector of correlations for the top behavioural variables (Fig. 2).

### Permutation framework

We used a permutation-based approach (Fig. S1) for choosing the optimal number of PCA components and estimating the Family Wise Error (FWE) corrected p-value on the canonical variates. The algorithm proceeds as follows:

1. For a given number of PCA components (e.g. *d = 5*), the reduction step is performed on *X* and *Y* resulting in the reduced data matrices *X*_*n*×*5*_ and *Y*_*n*×*5*_. Then, CCA is applied to this lower-dimensional space to compute the vector with “true” canonical correlations *q*_*5*×*1*_, with one value per CCA component.
2. The rows of data matrix *Y*_*n*×*5*_ are permuted to obtain 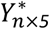 and then CCA is again computed using *X*_*n*×*5*_ and 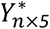 to obtain the corresponding vector of canonical correlations 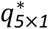. This procedure is repeated 10000 times, resulting in a matrix of canonical correlations 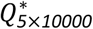.
3. For each row *i* = 1, …, 5 of 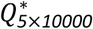, a p-value is computed by assessing the number of times the permuted canonical correlations in row *i* are equal or higher than the first “true” canonical correlation (as canonical correlations are ordered), i.e. the first element of *q*_*5*×*1*_ (equivalent to a maximum statistics approach). At the end of this procedure, a vector of p-values *P*_*5*×*1*_ is obtained, one per CCA component. This allows one to estimate the number of significant CCA components accounting for FWE (i.e. any CCA component with *P*_*FWE*_ < 0.05 is considered statistically significant). The p-value of the first CCA component (i.e. the first element of *P*_*5*×*1*_) is used to assess the optimal number of PCA components.
4. Steps 1-3 are repeated for the other PCA dimensionalities (*d* = 10,25,50,75,100,125,150,200).

The obtained p-values for each PCA dimensionality were corrected for multiple comparisons using *Bonferroni* correction (i.e. *α* = 0.05/9 = 0.0056), which means that only the PCA sets with *P*_*corr*_ ≤ 0.0056 are considered statistically significant. The optimal number of PCA components is chosen based on the lowest *P*_*corr*._

### Data availability

The datasets analysed during the current study are not publicly available because this was not foreseen when the ethical approval for the study was obtained, but are available from the corresponding author on reasonable request.

## Acknowledgments

The authors would like to acknowledge Prantik Kundu, Ameera Patel and Kirstie Whitaker for their contribution to MRI data pre-processing.

This work has been supported by a Wellcome Trust Strategic Award (reference: 095844) to Ian Goodyer, Raymond Dolan, Ed Bullmore, Peter Jones and Peter Fonagy which provides core funding for the Neuroscience in Psychiatry Network (NSPN.org.uk). Maria J. Rosa, Janaina Mourao-Miranda and Agoston Mihalik are funded by the Wellcome Trust under grant number WT102845/Z/13/Z. Joao M. Monteiro and Fabio S. Ferreira are funded by a PhD scholarship awarded by Fundacao para a Ciencia e a Tecnologia (SFRH/BD/88345/2012 and SFRH/BD/120640/2016, respectively). Michael Moutoussis is funded by the UCLH Biomedical Research Council. Rick A. Adams is supported by an MRC Skills Development Fellowship (MR/S007806/1). Petra E. Vértes was supported by the Medical Research Council grant number MR/K020706/1 and is a Fellow of MQ: Transforming Mental Health, grant number MQF17_24. František Váša was supported by the Gates Cambridge Trust. Peter Fonagy is in receipt of a National Institute for Health Research (NIHR) Senior Investigator Award (NF-SI-0514-10157). Peter Fonagy was in part supported by the NIHR Collaboration for Leadership in Applied Health Research and Care (CLAHRC) North Thames at Barts Health NHS Trust. The views expressed are those of the authors and not necessarily those of the NHS, the NIHR or the Department of Health.

## Contributions

A.M., F.S.F., M.J.R. and J.M-M. designed the study. M.M., E.T.B., P.F., I.M.G., P.B.J., R.D. and NSPN consortium collected the data. A.M., M.J.R., M.M., G.Z., J.M., R.R.G., P.E.V., M.G.K, F.V. and M.V. contributed to MRI and/or clinical data pre-processing. A.M., F.S.F., M.J.R. implemented the modelling analysis; J.M-M. supervised the modelling analysis. A.M., F.S.F., M.J.R., M.M. G.Z., L.P., R.A.A., J.M.M. discussed the interpretation of the results. A.M., F.S.F. and M.J.R. prepared figures and tables. All authors contributed to the manuscript. All authors approved the final version of the manuscript.

