## Supplementary material for "Brain-behaviour modes of covariation in healthy and clinically depressed young people"

### Methods

*Self-report questionnaires*

- Antisocial Behaviours Checklist (ABQ)[1] – self-report questionnaire for symptoms of antisocial behaviour based on DSM-IV conduct disorder items. The questionnaire was designed solely for the purpose of the NSPN project (11 items).
- Antisocial Process Screening Device (APSD)[2] – self-report scale measuring psychopathic traits and antisocial behaviour (20 items).
- Barratt Impulsive Scale (BIS)[3] – self-report questionnaire assessing personality and behavioural constructs of impulsiveness (30 items).
- Child and Adolescent Dispositions Scale (CADS)[4] – self-report measure of the three underlying dimensions of cognitive control of behaviour: pro-sociability, negative emotionality and daring (57 items).
- Child Trauma Questionnaire (CTQ)[5] – self-report inventory screening for histories of abuse and neglect, which covers five types of maltreatment: emotional, physical, and sexual abuse, and emotional and physical neglect (28 items).
- Drugs Alcohol and Self-Injury (DASI)[1] – self-report measure assessing the frequency of drug and alcohol use as well as the frequency, methods and motives of non-suicidal self-harm acts. The questionnaire was designed solely for the purpose of the NSPN project (16 items).
- Inventory of Callous-Unemotional Traits (ICU)[6] – self-report inventory of assessing 3 domains of callous and unemotional traits: callousness, uncaring, and unemotional (24 items).
- Kessler Psychological Distress Scale (K10)[7] – self-report measurement of psychological distress (10 items).
- Leyton Obsessional Inventory (LOI)[8] – self-report questionnaire measuring obsessional and anxiety symptoms (11 items).
- Moods and Feelings Questionnaire (MFQ)[9] – self-report questionnaire measuring depressive symptoms in the last 2 weeks (33 items).
- Revised Children's Manifest Anxiety Scale (RCMAS)[10] – self-report questionnaire measuring anxiety symptoms (28 items).
- Rosenberg Self-Esteem Scale (SES)[11] – self-report questionnaire measuring global self-esteem or feelings of self-worth and self-acceptance (10 items).
- Schizotypal Personality Questionnaire (SPQ)[12] – self-report scale measuring schizotypal personality traits (74 items).

- Wechsler Abbreviated Scale of Intelligence (WASI)[13] – matrix reasoning and vocabulary subsets of the Wechsler Abbreviated Scale of Intelligence (WASI) designed to assess fluid and crystallized intelligence, respectively (2 items).
- Warwick Edinburgh Mental Wellbeing Scale (WEMWBS)[14] – self-report instruments spanning the theoretical distribution of common mental symptoms and wellbeing (14 items).

*Socioeconomic deprivation index*

The socioeconomic deprivation index is a small-area model-based households in poverty estimate and reflects the proportion of deprived/poor households around the subject's residence (for full details see: https://www.ons.gov.uk/peoplepopulationandcommunity/personalandhouseholdfinances/incomeandwealth/bulletins/smallareamodelbasedhouseholdsinpovertyestimatesenglandandwales/financialyearending2014). The index was calculated by a searchable web page provided by the Office of National Statistics, UK (https://www.ons.gov.uk/), however, we note that the online tool is no longer available, and only a downloadable updated dataset can be found here: https://www.ons.gov.uk/peoplepopulationandcommunity/personalandhouseholdfinances/incomeandwealth/datasets/householdsinpovertyestimatesformiddlelayersuperoutputareasinenglandandwales.

*Brain visualization*

The glass brains were created using the BrainNet Viewer software[15]. In brief, BrainNet Viewer is a user-friendly MATLAB toolbox that was designed to visualize brain connectomes, as those in Figs. 3-4 and Figs. S4-5. The nodes were created using the coordinates of the centroids of the previously defined brain regions and the edges represent the brain connections found between the different regions, in which the red ones represent connections positively correlated with the canonical brain variates and the blue ones represent the ones negatively correlated with the canonical brain variates. The thickness of the edges is proportional to the absolute correlation value.

*Multiple hold-out framework*

We also used a multiple hold-out approach for choosing the optimal number of PCA components and estimating the Family Wise Error (FWE) corrected p-value on the canonical variates. This approach was implemented based on the framework proposed by Monteiro et. al[16] (Figure S2):

1. The data matrices $X_{n\times p}$ and $Y_{n\times q}$ are randomly split in two different sets of data, an optimisation set (80% of the total number of subjects), $X_{op}$ and $Y_{op}$, and a hold-out set (20% of the data), $X_{ho}$ and $Y_{ho}$ (Figure S2a).
2. For each PCA dimensionality $d$ ($d = 5,10,25,50,75,100,150,200$), the optimisation matrices, $X_{op}$ and $Y_{op}$, are randomly split 50 times into a training set (80% of the optimisation set), $X_{op_{tr}}$ and $Y_{op_{tr}}$, and a testing set (20% of the optimisation set), $X_{op_{te}}$ and $Y_{op_{te}}$. For each random split, the reduction step is performed on the

training set, $X_{op_{tr}}$ and $Y_{op_{tr}}$, and then CCA is applied to the reduced matrices, $X^d_{op_{tr}}$ and $Y^d_{op_{tr}}$, to obtain the combined PCA+CCA basis vectors $u_{tr}$ and $v_{tr}$. The test canonical correlations, $q_{te} \in \mathbb{R}^{d \times 1}$, are computed by projecting the testing set onto these basis vectors (Figure S2b). At the end of this procedure, for each PCA dimensionality *d*, a matrix of test canonical correlations ($Q_{te} \in \mathbb{R}^{d \times 50}$) is obtained, in which each row corresponds to a CCA component. The optimal number of PCA components is chosen based on the maximal average test canonical correlation of the first CCA components. In other words, the number of PCA components is chosen to maximise the test canonical CCA correlation (i.e. CCA correlation obtained when the test data was projected onto the PCA+CCA basis vectors estimated based on the train data).

3. PCA is applied to $X_{op}$ and $Y_{op}$ using the optimal number of components, $d^*$, resulting in the reduced data matrices $X_{d^*}$ and $Y_{d^*}$. $X_{d^*}$ and $Y_{d^*}$ are used to compute the combined PCA+CCA basis vectors, $u_{ho}$ and $v_{ho}$. The "true" hold-out canonical correlations, $q_{ho} \in \mathbb{R}^{d^* \times 1}$, are computed by projecting the hold-out set, $X_{ho}$ and $Y_{ho}$, onto these basis vectors (Figure S2c).
4. For assessing the statistical significance of the hold-out canonical correlations, we used permutation tests. First, the rows of $Y_{d^*}$, are permuted to obtain $Y^+_{d^*}$ then we applied CCA resulting in "permuted" combined PCA+CCA basis vectors $u^+_{ho}$ and $v^+_{ho}$. Third, we projected the hold-out set, $X_{ho}$and $Y_{ho}$, onto these basis vectors to obtain "permuted" hold-out canonical correlations $q^+_{ho} \in \mathbb{R}^{d^* \times 1}$. Next, we repeated the permutation procedure 10,000 times resulting in a matrix of "permuted" hold-out canonical correlations $Q^+_{ho} \in \mathbb{R}^{d^* \times 10000}$ (Fig. S6c). For each row of $Q^+_{ho}$ (representing a CCA component), a p-value is computed by assessing the number of times the permuted canonical correlations are equal or higher than the maximal "true" hold-out canonical correlation. We note that in contrast to the canonical correlations of the training set, the hold-out canonical correlations are not ordered. At the end of this procedure, a vector of p-values is obtained for each CCA component ($p \in \mathbb{R}^{d^* \times 1}$). This allows one to estimate the number of significant CCA components accounting for FWE (i.e. any CCA component with $p_{FWE} < 0.05$ is considered statistically significant).
5. Steps 1-4 are repeated 9 more times (10 different hold-out sets in total). The obtained p-values for each hold-out set are corrected for multiple comparisons using *Bonferroni* correction (i.e. $\alpha = 0.05/10 = 0.005$), which means that only the hold-out sets with $p_{corr} \leq 0.005$ are considered statistically significant. Finally, the best hold-out set is chosen based on the lowest $p_{corr}$.

### Tables

**Table S1:** CCA behaviour correlations computed by correlating the behavioural variables and the behavioural canonical variates of the first CCA mode, when using the permutation framework. Top 20 most positively and top 20 most negatively correlated items shown only.

| Clinical label | Questionnaire | Correlation |
|---|---|---|
| Male | Demographics | 0.68 |
| Age | Demographics | 0.38 |
| Do you enjoy doing things that are risky or dangerous? | CADS | 0.36 |
| I felt I had a number of good qualities | SES | 0.35 |
| During the last month, how often did you drink spirits? | DASI | 0.35 |
| Do you like TV, movies, comics, or electronic games with a lot of violence in them? | CADS | 0.34 |
| During the last month, how often did you drink beer or cider? | DASI | 0.34 |
| In the last 6 months, how often have you been drunk in the way described in Q7? | DASI | 0.33 |
| I was able to do things as well as most people | SES | 0.33 |
| Do you like rough games and sports? | CADS | 0.32 |
| Do you feel confident that you can handle life's challenges? | CADS | 0.28 |
| I've been able to make up my own mind about things | WEMWBS | 0.28 |
| Do you react with little or no emotion to both positive and negative things? | CADS | 0.28 |
| Are you calm and easy-going? | CADS | 0.28 |
| Are you proud of yourself? | CADS | 0.28 |
| I was satisfied with myself | SES | 0.28 |
| You tease or make fun of other people. | APSD | 0.27 |
| Are you energetic when you have a job to do? | CADS | 0.26 |
| I felt that I was as good as anyone else | SES | 0.26 |
| Raw Score (Vocabulary) | WASI | 0.26 |
| I sometimes blamed myself for things that weren't my fault | MFQ | -0.35 |
| At times, I thought I was no good at all | SES | -0.36 |
| Excluding the last month, have you tried to hurt yourself on purpose without trying to kill yourself in the last 12 months? | DASI | -0.36 |
| I did everything wrong | MFQ | -0.36 |
| I worried about what my parents would say to me | RCMAS | -0.36 |
| Excluding the last month, how have you tried to hurt yourself without trying to kill yourself in the last 12 months? | DASI | -0.37 |
| I thought about dying | MFQ | -0.37 |
| My feelings got hurt easily | RCMAS | -0.37 |
| Do you get upset easily? | CADS | -0.37 |
| In the last month, how have you tried to hurt yourself without trying to kill yourself? | DASI | -0.37 |
| I thought about killing myself | MFQ | -0.38 |
| I was afraid of a lot of things | RCMAS | -0.38 |

| | | |
|---|---|---|
| I worried what other people thought about me | RCMAS | -0.38 |
| Are you emotional? | CADS | -0.38 |
| Often I felt sick to my stomach | RCMAS | -0.39 |
| In the last month, why have you tried to hurt yourself without trying to kill yourself? | DASI | -0.39 |
| I thought my family would be better off without me | MFQ | -0.44 |
| I cried a lot | MFQ | -0.44 |
| I thought I looked ugly | MFQ | -0.48 |
| Female | Demographics | -0.68 |

**Table S2:** CCA connectivity correlations computed by correlating the brain connectivity variables and the brain canonical variates of the first CCA mode, computed using the permutation framework. Top 20 most positively and top 20 most negatively correlated brain connections shown only. **Functional networks:** Default Mode Network (DMN); Dorsal Attention Network (DAN); Frontoparietal Network (FPT); Limbic Network (LMB); Somatomotor Network (SMT); Ventral Attention Network (VAN); Subcortex (SBC); Visual Network (VIS). **Anatomical regions:** [1]Anterior Cingulate and Medial Prefrontal Cortex; [2]Auditory Association Cortex; [3]Basal Ganglia; [4]Dorsal Stream Visual Cortex; [5]Dorsolateral Prefrontal Cortex; [6]Early Auditory Cortex; [7]Early Visual Cortex; [8]Hippocampus; [9]Inferior Frontal Cortex; [10]Inferior Parietal Cortex; [11]Insular and Frontal Opercular Cortex; [12]Lateral Temporal Cortex; [13]MT+ Complex and Neighboring Visual Areas; [14]Medial Temporal Cortex; [15]Orbital and Polar Frontal Cortex; [16]Paracentral Lobular and Mid Cingulate Cortex; [17]Posterior Cingulate Cortex; [18]Posterior Opercular Cortex; [19]Premotor Cortex; [20]Primary Visual Cortex; [21]Somatosensory and Motor Cortex; [22]Superior Parietal Cortex; [23]Temporo-Parieto-Occipital Junction; [24]Thalamus; [25]Ventral Stream Visual Cortex.

| Label node A | Label node B | Correlation |
|---|---|---|
| Left-Insular-Granular-Complex (SMT)[11] | Left-Ventral-IntraParietal-Complex (DAN)[22] | 0.69 |
| Right-Lateral-Area-7P (DAN)[22] | Left-Insular-Granular-Complex (SMT)[11] | 0.68 |
| Left-Area-OP2-3/VS (SMT)[18] | Left-Ventral-IntraParietal-Complex (DAN)[22] | 0.68 |
| Left-Medial-Belt-Complex (SMT)[6] | Left-Ventral-IntraParietal-Complex (DAN)[22] | 0.67 |
| Right-Medial-Area-7A (DAN)[22] | Left-Insular-Granular-Complex (SMT)[11] | 0.67 |
| Right-Ventral-IntraParietal-Complex (DAN)[22] | Left-Insular-Granular-Complex (SMT)[11] | 0.67 |
| Right-Area-OP2-3/VS (SMT)[18] | Right-Ventral-IntraParietal-Complex (DAN)[22] | 0.67 |
| Right-Ventral-IntraParietal-Complex (DAN)[22] | Left-Area-OP2-3/VS (SMT)[18] | 0.66 |
| Right-Ventral-IntraParietal-Complex (DAN)[22] | Left-Medial-Belt-Complex (SMT)[6] | 0.66 |
| Right-Medial-Area-7P (FPT)[22] | Left-Frontal-Opercular-Area-3 (VAN)[11] | 0.66 |
| Right-Insular-Granular-Complex (SMT)[11] | Right-Ventral-IntraParietal-Complex | 0.66 |

| | | |
|---|---|---|
| | (DAN)[22] | |
| Right-Lateral-Area-7P (DAN)[22] | Left-Area-PFcm (VAN)[6] | 0.66 |
| Right-Medial-Belt-Complex (SMT)[6] | Right-Medial-Area-7A (DAN)[22] | 0.66 |
| Right-Ventral-IntraParietal-Complex (DAN)[22] | Left-RetroInsular-Cortex (SMT)[6] | 0.66 |
| Left-Area-PFcm (VAN)[6] | Left-Lateral-Area-7P (DAN)[22] | 0.66 |
| Right-Medial-Area-7P (FPT)[22] | Left-Area-PFcm (VAN)[6] | 0.65 |
| Right-Medial-Belt-Complex (SMT)[6] | Right-Ventral-IntraParietal-Complex (DAN)[22] | 0.65 |
| Right-Area-OP2-3/VS (SMT)[18] | Left-Ventral-IntraParietal-Complex (DAN)[22] | 0.65 |
| Right-Lateral-Area-7P (DAN)[22] | Left-Area-OP2-3/VS (SMT)[18] | 0.65 |
| Left-RetroInsular-Cortex (SMT)[6] | Left-Ventral-IntraParietal-Complex (DAN)[22] | 0.65 |
| Right-Area-10v (LMB)[1] | Left-Area-p32 (DMN)[1] | -0.09 |
| Left-Polar-10p (LMB)[15] | Left-Area-anterior-47r (DMN)[9] | -0.09 |
| Left-Polar-10p (LMB)[15] | Left-Area-8Av (DMN)[5] | -0.09 |
| Left-Polar-10p (LMB)[15] | Left-Area-10v (LMB)[1] | -0.10 |
| Right-Area-ventral-23-a+b (DMN)[17] | Left-Area-8Ad (DMN)[5] | -0.10 |
| Right-Area-31pd (DMN)[17] | Left-Parieto-Occipital-Sulcus-Area-2 (DMN)[17] | -0.10 |
| Right-Area-9-Middle (DMN)[1] | Left-Area-p32 (DMN)[1] | -0.10 |
| Right-Area-8Ad (DMN)[5] | Left-Area-PGs (DMN)[10] | -0.10 |
| Right-Area-ventral-23-a+b (DMN)[17] | Left-Area-7m (DMN)[17] | -0.10 |
| Right-Area-p32 (DMN)[1] | Left-Area-ventral-23-a+b (DMN)[17] | -0.10 |
| Left-Polar-10p (LMB)[15] | Left-Area-9-Posterior (DMN)[5] | -0.10 |
| Left-Area-10v (LMB)[1] | Left-Area-p32 (DMN)[1] | -0.10 |
| Left-Polar-10p (LMB)[15] | Left-Area-8C (FPT)[5] | -0.10 |
| Left-Polar-10p (LMB)[15] | Left-Area-8Ad (DMN)[5] | -0.11 |
| Right-Area-10v (LMB)[1] | Left-Polar-10p (LMB)[15] | -0.11 |
| Left-Area-PGs (DMN)[10] | Left-Polar-10p (LMB)[15] | -0.12 |
| Left-Area-PFm-Complex (DMN)[10] | Left-Polar-10p (LMB)[15] | -0.12 |
| Left-Area-posterior-10p (DMN)[15] | Left-Polar-10p (LMB)[15] | -0.12 |
| Left-Area-9-Middle (DMN)[1] | Left-Area-p32 (DMN)[1] | -0.12 |
| Right-Polar-10p (LMB)[15] | Left-Polar-10p (LMB)[15] | -0.13 |

**Table S3:** CCA behaviour correlations computed by correlating the behavioural variables and the behavioural canonical variates of the second CCA mode, when using the permutation framework. Top 20 most positively and top 20 most negatively correlated items shown only.

| Clinical label | Questionnaire | Correlation |
|---|---|---|
| I've been feeling cheerful | WEMWBS | 0.38 |
| I've been feeling interested in other people | WEMWBS | 0.38 |
| Are you cheerful? | CADS | 0.36 |
| Do you enjoy being with other people your age? | CADS | 0.33 |

| | | |
|---|---|---|
| I've been dealing with problems well | WEMWBS | 0.33 |
| I've been able to make up my own mind about things | WEMWBS | 0.30 |
| I've been feeling relaxed | WEMWBS | 0.30 |
| I've been feeling good about myself | WEMWBS | 0.28 |
| I've been feeling confident | WEMWBS | 0.27 |
| I've been feeling useful | WEMWBS | 0.27 |
| I am happy-go-lucky. | BIS | 0.27 |
| I've been feeling close to other people | WEMWBS | 0.26 |
| Do you feel confident that you can handle life's challenges? | CADS | 0.24 |
| I've been thinking clearly | WEMWBS | 0.24 |
| I've had energy to spare | WEMWBS | 0.23 |
| I've been interested in new things | WEMWBS | 0.23 |
| Are you enthusiastic about life? | CADS | 0.23 |
| Do you like things that are exciting and loud? | CADS | 0.22 |
| I've been feeling loved | WEMWBS | 0.22 |
| You lie easily and skilfully. | APSD | 0.21 |
| I hated myself | MFQ | -0.32 |
| I had bad dreams | RCMAS | -0.32 |
| I thought my family would be better off without me | MFQ | -0.33 |
| I worried a lot of the time | RCMAS | -0.33 |
| Someone molested me. | CTQ | -0.33 |
| I believe that I was sexually abused. | CTQ | -0.34 |
| Someone tried to make me do sexual things or watch sexual things. | CTQ | -0.34 |
| Have you ever been drunk? | DASI | -0.34 |
| I cried a lot | MFQ | -0.34 |
| In the last month, how have you tried to hurt yourself without trying to kill yourself? | DASI | -0.35 |
| I felt miserable or unhappy | MFQ | -0.35 |
| Someone tried to touch me in a sexual way, or tried to make me touch them. | CTQ | -0.35 |
| During the last month, how often did you drink spirits? | DASI | -0.35 |
| I thought there was nothing good for me in the future | MFQ | -0.36 |
| During the last 30 days, about how often did you feel so sad that nothing could cheer you up? | K10 | -0.36 |
| I thought about dying | MFQ | -0.37 |
| During the last 30 days, about how often did you feel depressed? | K10 | -0.37 |
| Age | Demographics | -0.38 |
| I thought that life was not worth living | MFQ | -0.46 |
| I thought about killing myself | MFQ | -0.47 |

**Table S4:** CCA connectivity correlations computed by correlating the brain connectivity variables and the brain canonical variates of the second CCA mode, computed using the permutation framework. Top 20 most positively and top 20 most negatively correlated brain connections shown only. **Functional networks:** Default Mode Network (DMN); Dorsal Attention Network (DAN); Frontoparietal Network (FPT); Limbic Network (LMB); Somatomotor Network (SMT); Ventral Attention Network (VAN); Subcortex (SBC); Visual Network (VIS). **Anatomical regions:** [1]Anterior Cingulate and Medial Prefrontal Cortex; [2]Auditory Association Cortex; [3]Basal Ganglia; [4]Dorsal Stream Visual Cortex; [5]DorsoLateral Prefrontal Cortex; [6]Early Auditory Cortex; [7]Early Visual Cortex; [8]Hippocampus; [9]Inferior Frontal Cortex; [10]Inferior Parietal Cortex; [11]Insular and Frontal Opercular Cortex; [12]Lateral Temporal Cortex; [13]MT+ Complex and Neighboring Visual Areas; [14]Medial Temporal Cortex; [15]Orbital and Polar Frontal Cortex; [16]Paracentral Lobular and Mid Cingulate Cortex; [17]Posterior Cingulate Cortex; [18]Posterior Opercular Cortex; [19]Premotor Cortex; [20]Primary Visual Cortex; [21]Somatosensory and Motor Cortex; [22]Superior Parietal Cortex; [23]Temporo-Parieto-Occipital Junction; [24]Thalamus; [25]Ventral Stream Visual Cortex.

| Label node A | Label node B | Correlation |
|---|---|---|
| Left-Area-8Ad (DMN)[5] | Left-Thalamus-Proper (SBC)[24] | 0.38 |
| Right-Area-dorsal-23-a+b (DMN)[17] | Left-Superior-Frontal-Language-Area (DMN)[5] | 0.37 |
| Right-Area-7m (DMN)[17] | Left-Thalamus-Proper (SBC)[24] | 0.36 |
| Left-Area-dorsal-23-a+b (DMN)[17] | Left-Superior-Frontal-Language-Area (DMN)[5] | 0.35 |
| Left-Area-posterior-24 (DMN)[1] | Left-Area-8Ad (DMN)[5] | 0.35 |
| Left-Area-PGs (DMN)[10] | Left-Thalamus-Proper (SBC)[24] | 0.35 |
| Left-Area-7m (DMN)[17] | Left-Thalamus-Proper (SBC)[24] | 0.35 |
| Right-Area-8Ad (DMN)[5] | Left-Thalamus-Proper (SBC)[24] | 0.34 |
| Right-Area-31p-ventral (DMN)[17] | Left-Superior-Frontal-Language-Area (DMN)[5] | 0.34 |
| Left-Area-posterior-24 (DMN)[1] | Left-Area-dorsal-23-a+b (DMN)[17] | 0.34 |
| Left-Area-posterior-24 (DMN)[1] | Left-Area-8Av (DMN)[5] | 0.34 |
| Right-Area-dorsal-23-a+b (DMN)[17] | Left-Area-posterior-24 (DMN)[1] | 0.34 |
| Left-Area-posterior-24 (DMN)[1] | Left-Area-PGs (DMN)[10] | 0.33 |
| Right-Area-5m (SMT)[16] | Right-Area-dorsal-23-a+b (DMN)[17] | 0.32 |
| Right-Area-8Ad (DMN)[5] | Left-Area-posterior-24 (DMN)[1] | 0.32 |
| Left-Area-TE1-posterior (DMN)[12] | Left-Area-9-Middle (DMN)[1] | 0.32 |
| Left-Superior-Frontal-Language-Area (DMN)[5] | Left-Thalamus-Proper (SBC)[24] | 0.32 |
| Left-Area-9-Middle (DMN)[1] | Left-Thalamus-Proper (SBC)[24] | 0.32 |
| Left-Area-TE1-posterior (DMN)[12] | Left-Area-9-anterior (DMN)[5] | 0.32 |
| Right-Area-dorsal-23-a+b (DMN)[17] | Left-Area-6m-anterior (VAN)[16] | 0.32 |
| Right-Area-PF-opercular (VAN)[10] | Right-Area-5m (SMT)[16] | -0.27 |
| Right-Area-PFt (DAN)[10] | Left-Middle-Temporal-Area (VIS)[13] | -0.27 |
| Right-Area-PFt (DAN)[10] | Left-Area-Lateral-Occipital-1 (VIS)[13] | -0.28 |
| Right-RetroInsular-Cortex (SMT)[6] | Left-Area-PF-opercular (VAN)[10] | -0.28 |
| Right-Area-PFt (DAN)[10] | Left-Seventh-Visual-Area (VIS)[4] | -0.28 |
| Right-Area-PFcm (SMT)[6] | Left-Area-6m-anterior (VAN)[16] | -0.28 |

| | | |
|---|---|---|
| Right-ParaBelt-Complex (SMT)[6] | Left-Frontal-Opercular-Area-2 (SMT)[11] | -0.28 |
| Right-Area-PFt (DAN)[10] | Left-Area-V3CD (VIS)[13] | -0.28 |
| Right-Auditory-4-Complex (SMT)[2] | Left-Frontal-Opercular-Area-2 (SMT)[11] | -0.28 |
| Right-Frontal-Opercular-Area-4 (VAN)[11] | Left-Area-Posterior-Insular-1 (VAN)[11] | -0.28 |
| Right-Area-PFt (DAN)[10] | Right-Middle-Temporal-Area (VIS)[13] | -0.28 |
| Right-Area-PFcm (SMT)[6] | Left-Area-Frontal-Opercular-5 (VAN)[11] | -0.29 |
| Right-Area-V4t (VIS)[13] | Right-Area-PFt (DAN)[10] | -0.29 |
| Right-Area-Lateral-Occipital-3 (VIS)[13] | Right-Area-PFt (DAN)[10] | -0.29 |
| Right-Frontal-Opercular-Area-4 (VAN)[11] | Right-Area-PFcm (SMT)[6] | -0.30 |
| Right-RetroInsular-Cortex (SMT)[6] | Left-Frontal-Opercular-Area-2 (SMT)[11] | -0.30 |
| Right-Area-PF-opercular (VAN)[10] | Left-Area-6m-anterior (VAN)[16] | -0.30 |
| Right-Area-PFt (DAN)[10] | Right-Area-Lateral-Occipital-2 (VIS)[13] | -0.31 |
| Right-Area-5m-ventral (VAN)[16] | Left-Area-Frontal-Opercular-5 (VAN)[11] | -0.31 |
| Right-Area-PFt (DAN)[10] | Right-Area-Lateral-Occipital-1 (VIS)[13] | -0.31 |

**Table S5:** CCA connectivity correlations computed by correlating the behavioural variables and the behavioural canonical variates of the first CCA mode, when using the multiple hold-out framework. Top 20 most positively and top 20 most negatively correlated items shown only.

| Clinical label | Questionnaire | Correlation |
|---|---|---|
| Age | Demographics | 0.53 |
| During the last month, how often did you drink spirits? | DASI | 0.48 |
| In the last 6 months, how often have you been drunk in the way described in Q7? | DASI | 0.47 |
| Have you ever been drunk? | DASI | 0.47 |
| Male | Demographics | 0.42 |
| During the last month, how often did you drink beer or cider? | DASI | 0.41 |
| During the last month, how often did you smoke a cigarette/s? | DASI | 0.37 |
| During the last month, on the days you smoked, on average how many cigarettes did you smoke per day? | DASI | 0.36 |
| You engage in illegal activities. | APSD | 0.35 |
| Do you react with little or no emotion to both positive and negative things? | CADS | 0.32 |
| Do you like TV, movies, comics, or electronic games with a lot of violence in them? | CADS | 0.31 |
| Do you enjoy doing things that are risky or dangerous? | CADS | 0.31 |
| I like to think about complex problems. | BIS | 0.29 |
| I seem very cold and uncaring to others. | ICU | 0.28 |
| You do risky or dangerous things. | APSD | 0.27 |
| I do not show my emotions to others. | ICU | 0.26 |
| During the last month, how often did you drink wine? | DASI | 0.25 |
| During the last month, how often did you take/use cannabis? | DASI | 0.25 |

| | | |
|---|---|---|
| Are you brave? | CADS | 0.24 |
| Are you daring and adventurous? | CADS | 0.23 |
| I sometimes jump quickly from one topic to another when speaking. | SPQ | -0.20 |
| I worried what other people thought about me | RCMAS | -0.20 |
| Do you want everyone to follow the rules, including yourself? | CADS | -0.20 |
| I had the best family in the world. | CTQ | -0.20 |
| I often ramble on too much when speaking. | SPQ | -0.21 |
| I thought my family would be better off without me | MFQ | -0.21 |
| I am concerned about the feelings of others. | ICU | -0.21 |
| Do you sometimes feel that other people are watching you? | SPQ | -0.22 |
| Do you sometimes feel that people are talking about you? | SPQ | -0.22 |
| When you see other people talking to each other, do you often wonder if they are talking about you? | SPQ | -0.22 |
| My feelings got hurt easily when I was fussed at | RCMAS | -0.23 |
| Do you like things to stay the same and not change? | CADS | -0.24 |
| My feelings got hurt easily | RCMAS | -0.24 |
| Do you get upset easily? | CADS | -0.24 |
| I thought I looked ugly | MFQ | -0.25 |
| Are you emotional? | CADS | -0.27 |
| Are you easily embarrassed? | CADS | -0.29 |
| I worried about what my parents would say to me | RCMAS | -0.32 |
| Would you feel guilty if you did something that broke the law? | CADS | -0.36 |
| Female | Demographics | -0.42 |

**Table S6:** CCA connectivity correlations computed by correlating the brain connectivity variables and the brain canonical variates of the first CCA mode, when using the multiple hold-out framework. Top 20 most positively and top 20 most negatively correlated brain connections shown only. **Functional networks:** Default Mode Network (DMN); Dorsal Attention Network (DAN); Frontoparietal Network (FPT); Limbic Network (LMB); Somatomotor Network (SMT); Ventral Attention Network (VAN); Subcortex (SBC); Visual Network (VIS). **Anatomical regions:** [1]Anterior Cingulate and Medial Prefrontal Cortex; [2]Auditory Association Cortex; [3]Basal Ganglia; [4]Dorsal Stream Visual Cortex; [5]DorsoLateral Prefrontal Cortex; [6]Early Auditory Cortex; [7]Early Visual Cortex; [8]Hippocampus; [9]Inferior Frontal Cortex; [10]Inferior Parietal Cortex; [11]Insular and Frontal Opercular Cortex; [12]Lateral Temporal Cortex; [13]MT+ Complex and Neighboring Visual Areas; [14]Medial Temporal Cortex; [15]Orbital and Polar Frontal Cortex; [16]Paracentral Lobular and Mid Cingulate Cortex; [17]Posterior Cingulate Cortex; [18]Posterior Opercular Cortex; [19]Premotor Cortex; [20]Primary Visual Cortex; [21]Somatosensory and Motor Cortex; [22]Superior Parietal Cortex; [23]Temporo-Parieto-Occipital Junction; [24]Thalamus; [25]Ventral Stream Visual Cortex.

| Label node A | Label node B | Correlation |
|---|---|---|
| Right-Insular-Granular-Complex (SMT)[11] | Right-Third-Visual-Area (VIS)[7] | 0.71 |

| | | |
|---|---|---|
| Right-Medial-Area-7A (DAN)[22] | Left-Insular-Granular-Complex (SMT)[11] | 0.71 |
| Right-Posterior-Insular-Area-2 (VAN)[11] | Left-Area-5m (SMT)[16] | 0.71 |
| Left-Insular-Granular-Complex (SMT)[11] | Left-Sixth-Visual-Area (VIS)[4] | 0.71 |
| Left-Insular-Granular-Complex (SMT)[11] | Left-Area-5m (SMT)[16] | 0.71 |
| Right-PreCuneus-Visual-Area (DAN)[17] | Left-Insular-Granular-Complex (SMT)[11] | 0.71 |
| Right-PreCuneus-Visual-Area (DAN)[17] | Left-Area-OP2-3/VS (SMT)[18] | 0.71 |
| Right-Insular-Granular-Complex (SMT)[11] | Left-Area-5m (SMT)[16] | 0.70 |
| Right-Area-OP2-3/VS (SMT)[18] | Left-Area-5m (SMT)[16] | 0.70 |
| Right-Insular-Granular-Complex (SMT)[11] | Right-Second-Visual-Area (VIS)[7] | 0.70 |
| Right-Area-23c (VAN)[16] | Left-Area-OP2-3/VS (SMT)[18] | 0.70 |
| Left-Area-OP2-3/VS (SMT)[18] | Left-Area-5m (SMT)[16] | 0.70 |
| Right-Insular-Granular-Complex (SMT)[11] | Left-Third-Visual-Area (VIS)[7] | 0.70 |
| Left-Insular-Granular-Complex (SMT)[11] | Left-PreCuneus-Visual-Area (DMN)[17] | 0.70 |
| Right-Insular-Granular-Complex (SMT)[11] | Left-Fourth-Visual-Area (VIS)[7] | 0.70 |
| Right-Second-Visual-Area (VIS)[7] | Left-Insular-Granular-Complex (SMT)[11] | 0.70 |
| Right-Medial-Area-7A (DAN)[22] | Left-Area-OP2-3/VS (SMT)[18] | 0.69 |
| Left-Area-OP2-3/VS (SMT)[18] | Left-Sixth-Visual-Area (VIS)[4] | 0.69 |
| Right-Area-V3B (VIS)[4] | Left-Insular-Granular-Complex (SMT)[11] | 0.69 |
| Right-Area-5m-ventral (VAN)[16] | Left-Area-OP2-3/VS (SMT)[18] | 0.69 |
| Right-Area-23d (DMN)[17] | Left-Area-31pd (DMN)[17] | -0.13 |
| Left-Polar-10p (LMB)[15] | Left-Area-10v (LMB)[1] | -0.13 |
| Right-Area-10v (LMB)[1] | Left-Parieto-Occipital-Sulcus-Area-2 (DMN)[17] | -0.13 |
| Left-Area-posterior-10p (DMN)[15] | Left-Polar-10p (LMB)[15] | -0.13 |
| Left-Area-ventral-23-a+b (DMN)[17] | Left-Area-23d (DMN)[17] | -0.13 |
| Right-Area-10v (LMB)[1] | Left-Area-23d (DMN)[17] | -0.13 |
| Left-Area-PGs (DMN)[10] | Left-Polar-10p (LMB)[15] | -0.13 |
| Right-Area-23d (DMN)[17] | Left-Area-31p-ventral (DMN)[17] | -0.13 |
| Left-Area-10v (LMB)[1] | Left-Area-23d (DMN)[17] | -0.14 |
| Right-RetroSplenial-Complex (DMN)[17] | Left-Area-31pd (DMN)[17] | -0.14 |
| Right-Area-10v (LMB)[1] | Left-Polar-10p (LMB)[15] | -0.14 |
| Left-Area-31p-ventral (DMN)[17] | Left-Parieto-Occipital-Sulcus-Area-2 (DMN)[17] | -0.14 |
| Left-Area-10v (LMB)[1] | Left-Parieto-Occipital-Sulcus-Area-2 (DMN)[17] | -0.14 |
| Right-Area-8Ad (DMN)[5] | Left-Area-PGs (DMN)[10] | -0.15 |
| Left-Area-dorsal-23-a+b (DMN)[17] | Left-Area-23d (DMN)[17] | -0.15 |
| Right-Area-10v (LMB)[1] | Left-Area-p32 (DMN)[1] | -0.15 |
| Left-Area-31p-ventral (DMN)[17] | Left-Area-23d (DMN)[17] | -0.15 |
| Left-Area-31pd (DMN)[17] | Left-Area-23d (DMN)[17] | -0.16 |
| Left-Area-31pd (DMN)[17] | Left-Parieto-Occipital-Sulcus-Area-2 (DMN)[17] | -0.17 |
| Right-Polar-10p (LMB)[15] | Left-Polar-10p (LMB)[15] | -0.17 |

**Table S7:** Atlas with region MNI coordinates (x, y, z), region labels, anatomical and functional assignments.

| X | Y | Z | Label | Region | Networks | Networks_Short |
|---|---|---|---|---|---|---|
| -12 | -20 | 5 | Left-Thalamus-Proper | Thalamus | Subcortical Network | SBC |
| -15 | 9 | 7 | Left-Caudate | Basal Ganglia | Subcortical Network | SBC |
| -26 | 0 | -2 | Left-Putamen | Basal Ganglia | Subcortical Network | SBC |
| -26 | -24 | -15 | Left-Hippocampus | Hippocampus | Subcortical Network | SBC |
| -24 | -7 | -21 | Left-Amygdala | Basal Ganglia | Subcortical Network | SBC |
| -10 | 11 | -9 | Left-Accumbens-area | Basal Ganglia | Subcortical Network | SBC |
| 12 | -19 | 6 | Right-Thalamus-Proper | Thalamus | Subcortical Network | SBC |
| 16 | 9 | 8 | Right-Caudate | Basal Ganglia | Subcortical Network | SBC |
| 27 | 1 | -2 | Right-Putamen | Basal Ganglia | Subcortical Network | SBC |
| 28 | -23 | -15 | Right-Hippocampus | Hippocampus | Subcortical Network | SBC |
| 25 | -6 | -21 | Right-Amygdala | Basal Ganglia | Subcortical Network | SBC |
| 10 | 10 | -9 | Right-Accumbens-area | Basal Ganglia | Subcortical Network | SBC |
| -12 | -83 | 1 | Left-Primary-Visual-Cortex | Primary Visual Cortex | Visual Network | VIS |
| -42 | -67 | 4 | Left-Medial-Superior-Temporal-Area | MT+ Complex and Neighboring Visual Areas | Dorsal Attention Network | DAN |
| -14 | -79 | 28 | Left-Sixth-Visual-Area | Dorsal Strem Visual Cortex | Visual Network | VIS |
| -12 | -80 | 2 | Left-Second-Visual-Area | Early Visual Cortex | Visual Network | VIS |
| -18 | -85 | 5 | Left-Third-Visual-Area | Early Visual Cortex | Visual Network | VIS |
| -30 | -83 | -4 | Left-Fourth-Visual-Area | Early Visual Cortex | Visual Network | VIS |
| -33 | -71 | -15 | Left-Eighth-Visual-Area | Ventral Stream Visual Cortex | Visual Network | VIS |
| -28 | -20 | 54 | Left-Primary-Motor-Cortex | Somatosensory and Motor Cortex | Somatomotor Network | SMT |
| -38 | -23 | 51 | Left-Primary-Sensory-Cortex | Somatosensory and Motor Cortex | Somatomotor Network | SMT |
| -40 | -5 | 48 | Left-Frontal-Eye-Fields | Premotor Cortex | Dorsal Attention Network | DAN |
| -48 | -1 | 38 | Left-Premotor-Eye-Field | Premotor Cortex | Dorsal Attention Network | DAN |
| -48 | -1 | 46 | Left-Area-55b | Premotor Cortex | Somatomotor Network | SMT |
| -16 | -89 | 24 | Left-Area-V3A | Dorsal Strem Visual Cortex | Visual Network | VIS |
| -6 | -40 | 17 | Left-RetroSplenial-Complex | Posterior Cingulate Cortex | Default Mode Network | DMN |
| -10 | -70 | 34 | Left-Parieto-Occipital-Sulcus-Area-2 | Posterior Cingulate Cortex | Default Mode Network | DMN |

| | | | | | | |
|---|---|---|---|---|---|---|
| -24 | -82 | 26 | Left-Seventh-Visual-Area | Dorsal Strem Visual Cortex | Visual Network | VIS |
| -23 | -73 | 33 | Left-IntraParietal-Sulcus-Area-1 | Dorsal Strem Visual Cortex | Dorsal Attention Network | DAN |
| -41 | -56 | -20 | Left-Fusiform-Face-Complex | Ventral Stream Visual Cortex | Visual Network | VIS |
| -27 | -80 | 16 | Left-Area-V3B | Dorsal Strem Visual Cortex | Visual Network | VIS |
| -39 | -84 | 2 | Left-Area-Lateral-Occipital-1 | MT+ Complex and Neighboring Visual Areas | Visual Network | VIS |
| -43 | -82 | -7 | Left-Area-Lateral-Occipital-2 | MT+ Complex and Neighboring Visual Areas | Visual Network | VIS |
| -41 | -80 | -14 | Left-Posterior-InferoTemporal-Complex | Ventral Stream Visual Cortex | Visual Network | VIS |
| -45 | -73 | 6 | Left-Middle-Temporal-Area | MT+ Complex and Neighboring Visual Areas | Visual Network | VIS |
| -44 | -24 | 9 | Left-Primary-Auditory-Cortex | Early Auditory Cortex | Somatomotor Network | SMT |
| -57 | -45 | 21 | Left-PeriSylvian-Language-Area | Temporo-Parieto-Occipital Junction | Ventral Attention Network | VAN |
| -8 | 17 | 61 | Left-Superior-Frontal-Language-Area | DorsoLateral Prefrontal Cortex | Default Mode Network | DMN |
| -7 | -50 | 48 | Left-PreCuneus-Visual-Area | Posterior Cingulate Cortex | Default Mode Network | DMN |
| -60 | -47 | 14 | Left-Superior-Temporal-Visual-Area | Temporo-Parieto-Occipital Junction | Default Mode Network | DMN |
| -7 | -66 | 48 | Left-Medial-Area-7P | Superior Parietal Cortex | Frontoparietal Network | FPT |
| -5 | -62 | 33 | Left-Area-7m | Posterior Cingulate Cortex | Default Mode Network | DMN |
| -11 | -59 | 13 | Left-Parieto-Occipital-Sulcus-Area-1 | Posterior Cingulate Cortex | Default Mode Network | DMN |
| -3 | -20 | 37 | Left-Area-23d | Posterior Cingulate Cortex | Default Mode Network | DMN |
| -5 | -56 | 18 | Left-Area-ventral-23-a+b | Posterior Cingulate Cortex | Default Mode Network | DMN |
| -3 | -40 | 30 | Left-Area-dorsal-23-a+b | Posterior Cingulate Cortex | Default Mode Network | DMN |
| -8 | -45 | 32 | Left-Area-31p-ventral | Posterior Cingulate Cortex | Default Mode Network | DMN |
| -7 | -39 | 60 | Left-Area-5m | Paracentral Lobular and Mid Cingulate Cortex | Somatomotor Network | SMT |
| -15 | -36 | 48 | Left-Area-5m-ventral | Paracentral Lobular and Mid Cingulate Cortex | Ventral Attention Network | VAN |
| -11 | -28 | 42 | Left-Area-23c | Paracentral Lobular and Mid Cingulate Cortex | Ventral Attention Network | VAN |
| -13 | -46 | 69 | Left-Area-5L | Paracentral Lobular and Mid Cingulate Cortex | Somatomotor Network | SMT |
| -8 | -18 | 49 | Left-Dorsal-Area-24d | Paracentral Lobular and Mid Cingulate Cortex | Somatomotor Network | SMT |
| -10 | -2 | 43 | Left-Ventral-Area-24d | Paracentral Lobular and Mid Cingulate Cortex | Somatomotor Network | SMT |
| -19 | -53 | 64 | Left-Lateral-Area-7A | Superior Parietal Cortex | Dorsal Attention Network | DAN |
| -7 | 3 | 56 | Left-Supplementary-and-Cingulate-Eye-Field | Paracentral Lobular and Mid Cingulate Cortex | Ventral Attention Network | VAN |
| -17 | 4 | 66 | Left-Area-6m-anterior | Paracentral Lobular and Mid Cingulate Cortex | Ventral Attention Network | VAN |

| | | | | | | |
|---|---|---|---|---|---|---|
| -9 | -58 | 59 | Left-Medial-Area-7A | Superior Parietal Cortex | Dorsal Attention Network | DAN |
| -12 | -72 | 51 | Left-Lateral-Area-7P | Superior Parietal Cortex | Dorsal Attention Network | DAN |
| -34 | -50 | 59 | Left-Area-7PC | Superior Parietal Cortex | Dorsal Attention Network | DAN |
| -29 | -56 | 53 | Left-Area-Lateral-IntraParietal-ventral | Superior Parietal Cortex | Dorsal Attention Network | DAN |
| -21 | -64 | 60 | Left-Ventral-IntraParietal-Complex | Superior Parietal Cortex | Dorsal Attention Network | DAN |
| -22 | -64 | 42 | Left-Medial-IntraParietal-Area | Superior Parietal Cortex | Dorsal Attention Network | DAN |
| -44 | -26 | 54 | Left-Area-1 | Somatosensory and Motor Cortex | Somatomotor Network | SMT |
| -37 | -34 | 51 | Left-Area-2 | Somatosensory and Motor Cortex | Somatomotor Network | SMT |
| -34 | -21 | 41 | Left-Area-3a | Somatosensory and Motor Cortex | Somatomotor Network | SMT |
| -34 | -14 | 64 | Left-Dorsal-area-6 | Premotor Cortex | Somatomotor Network | SMT |
| -12 | -15 | 67 | Left-Area-6mp | Paracentral Lobular and Mid Cingulate Cortex | Somatomotor Network | SMT |
| -58 | 2 | 30 | Left-Ventral-Area-6 | Premotor Cortex | Somatomotor Network | SMT |
| -4 | -1 | 38 | Left-Area-Posterior-24-prime | Anterior Cingulate and Medial Prefrontal Cortex | Ventral Attention Network | VAN |
| -4 | 12 | 28 | Left-Area-33-prime | Anterior Cingulate and Medial Prefrontal Cortex | Frontoparietal Network | FPT |
| -5 | 19 | 29 | Left-Anterior-24-prime | Anterior Cingulate and Medial Prefrontal Cortex | Ventral Attention Network | VAN |
| -10 | 15 | 37 | Left-Area-p32-prime | Anterior Cingulate and Medial Prefrontal Cortex | Ventral Attention Network | VAN |
| -5 | 39 | -1 | Left-Area-a24 | Anterior Cingulate and Medial Prefrontal Cortex | Default Mode Network | DMN |
| -9 | 39 | 23 | Left-Area-dorsal-32 | Anterior Cingulate and Medial Prefrontal Cortex | Default Mode Network | DMN |
| -6 | 31 | 43 | Left-Area-8BM | Anterior Cingulate and Medial Prefrontal Cortex | Frontoparietal Network | FPT |
| -11 | 46 | -1 | Left-Area-p32 | Anterior Cingulate and Medial Prefrontal Cortex | Default Mode Network | DMN |
| -7 | 50 | -8 | Left-Area-10r | Anterior Cingulate and Medial Prefrontal Cortex | Default Mode Network | DMN |
| -35 | 31 | -15 | Left-Area-47m | Orbital and Polar Frontal Cortex | Default Mode Network | DMN |
| -38 | 17 | 49 | Left-Area-8Av | DorsoLateral Prefrontal Cortex | Default Mode Network | DMN |
| -23 | 26 | 42 | Left-Area-8Ad | DorsoLateral Prefrontal Cortex | Default Mode Network | DMN |
| -7 | 52 | 24 | Left-Area-9-Middle | Anterior Cingulate and Medial Prefrontal Cortex | Default Mode Network | DMN |
| -12 | 36 | 51 | Left-Area-8B-Lateral | DorsoLateral Prefrontal Cortex | Default Mode Network | DMN |
| -17 | 45 | 38 | Left-Area-9-Posterior | DorsoLateral Prefrontal Cortex | Default Mode Network | DMN |
| -11 | 63 | 9 | Left-Area-10d | Orbital and Polar Frontal Cortex | Default Mode Network | DMN |
| -41 | 17 | 36 | Left-Area-8C | DorsoLateral Prefrontal Cortex | Frontoparietal Network | FPT |

| | | | | | | |
|---|---|---|---|---|---|---|
| -51 | 15 | 12 | Left-Area-44 | Inferior Frontal Cortex | Default Mode Network | DMN |
| -48 | 26 | 4 | Left-Area-45 | Inferior Frontal Cortex | Default Mode Network | DMN |
| -44 | 31 | -11 | Left-Area-47l-(47-lateral) | Inferior Frontal Cortex | Default Mode Network | DMN |
| -39 | 47 | -10 | Left-Area-anterior-47r | Inferior Frontal Cortex | Default Mode Network | DMN |
| -52 | 6 | 17 | Left-Rostral-Area-6 | Premotor Cortex | Ventral Attention Network | VAN |
| -41 | 12 | 24 | Left-Area-IFJa | Inferior Frontal Cortex | Frontoparietal Network | FPT |
| -39 | 3 | 28 | Left-Area-IFJp | Inferior Frontal Cortex | Dorsal Attention Network | DAN |
| -47 | 21 | 20 | Left-Area-IFSp | Inferior Frontal Cortex | Frontoparietal Network | FPT |
| -45 | 31 | 11 | Left-Area-IFSa | Inferior Frontal Cortex | Frontoparietal Network | FPT |
| -43 | 29 | 28 | Left-Area-posterior-9-46v | DorsoLateral Prefrontal Cortex | Frontoparietal Network | FPT |
| -37 | 37 | 28 | Left-Area-46 | DorsoLateral Prefrontal Cortex | Frontoparietal Network | FPT |
| -37 | 48 | 9 | Left-Area-anterior-9-46v | DorsoLateral Prefrontal Cortex | Frontoparietal Network | FPT |
| -29 | 43 | 21 | Left-Area-9-46d | DorsoLateral Prefrontal Cortex | Frontoparietal Network | FPT |
| -19 | 54 | 25 | Left-Area-9-anterior | DorsoLateral Prefrontal Cortex | Default Mode Network | DMN |
| -4 | 51 | -18 | Left-Area-10v | Anterior Cingulate and Medial Prefrontal Cortex | Limbic Network | LMB |
| -25 | 56 | -6 | Left-Area-anterior-10p | Orbital and Polar Frontal Cortex | Default Mode Network | DMN |
| -12 | 58 | -16 | Left-Polar-10p | Orbital and Polar Frontal Cortex | Limbic Network | LMB |
| -24 | 46 | -14 | Left-Area-11l | Orbital and Polar Frontal Cortex | Frontoparietal Network | FPT |
| -23 | 26 | -19 | Left-Area-13l | Orbital and Polar Frontal Cortex | Limbic Network | LMB |
| -32 | 21 | -18 | Left-Area-47s | Orbital and Polar Frontal Cortex | Default Mode Network | DMN |
| -29 | -55 | 41 | Left-Area-Lateral-IntraParietal-dorsal | Superior Parietal Cortex | Dorsal Attention Network | DAN |
| -25 | -4 | 53 | Left-Area-6-anterior | Premotor Cortex | Dorsal Attention Network | DAN |
| -32 | 7 | 53 | Left-Inferior-6-8-Transitional-Area | DorsoLateral Prefrontal Cortex | Frontoparietal Network | FPT |
| -22 | 22 | 55 | Left-Superior-6-8-Transitional-Area | DorsoLateral Prefrontal Cortex | Default Mode Network | DMN |
| -56 | -1 | 9 | Left-Area-43 | Posterior Opercular Cortex | Ventral Attention Network | VAN |
| -58 | -13 | 14 | Left-Area-OP4/PV | Posterior Opercular Cortex | Somatomotor Network | SMT |
| -47 | -22 | 17 | Left-Area-OP1/SII | Posterior Opercular Cortex | Somatomotor Network | SMT |
| -40 | -16 | 17 | Left-Area-OP2-3/VS | Posterior Opercular Cortex | Somatomotor Network | SMT |
| -39 | -22 | -2 | Left-Area-52 | Early Auditory Cortex | Somatomotor Network | SMT |

| | | | | | | |
|---|---|---|---|---|---|---|
| -40 | -35 | 17 | Left-RetroInsular-Cortex | Early Auditory Cortex | Somatomotor Network | SMT |
| -51 | -32 | 19 | Left-Area-PFcm | Early Auditory Cortex | Ventral Attention Network | VAN |
| -39 | -4 | -2 | Left-Posterior-Insular-Area-2 | Insular and Frontal Opercular Cortex | Ventral Attention Network | VAN |
| -51 | 0 | -8 | Left-Area-TA2 | Auditory Association Cortex | Somatomotor Network | SMT |
| -40 | 12 | 6 | Left-Frontal-Opercular-Area-4 | Insular and Frontal Opercular Cortex | Ventral Attention Network | VAN |
| -36 | 9 | 1 | Left-Middle-Insular-Area | Insular and Frontal Opercular Cortex | Ventral Attention Network | VAN |
| -30 | 5 | -18 | Left-Pirform-Cortex | Insular and Frontal Opercular Cortex | Limbic Network | LMB |
| -31 | 23 | -3 | Left-Anterior-Ventral-Insular-Area | Insular and Frontal Opercular Cortex | Frontoparietal Network | FPT |
| -34 | 13 | -12 | Left-Anterior-Agranular-Insula-Complex | Insular and Frontal Opercular Cortex | Default Mode Network | DMN |
| -50 | 3 | 4 | Left-Frontal-Opercular-Area-1 | Posterior Opercular Cortex | Ventral Attention Network | VAN |
| -35 | 3 | 12 | Left-Frontal-Opercular-Area-3 | Insular and Frontal Opercular Cortex | Ventral Attention Network | VAN |
| -42 | -4 | 13 | Left-Frontal-Opercular-Area-2 | Insular and Frontal Opercular Cortex | Somatomotor Network | SMT |
| -50 | -30 | 36 | Left-Area-PFt | Inferior Parietal Cortex | Dorsal Attention Network | DAN |
| -36 | -42 | 41 | Left-Anterior-IntraParietal-Area | Superior Parietal Cortex | Dorsal Attention Network | DAN |
| -17 | -35 | -11 | Left-PreSubiculum | Medial Temporal Cortex | Default Mode Network | DMN |
| -21 | -55 | 2 | Left-ProStriate-Area | Posterior Cingulate Cortex | Visual Network | VIS |
| -51 | 9 | -18 | Left-Area-STGa | Auditory Association Cortex | Default Mode Network | DMN |
| -54 | -25 | 5 | Left-ParaBelt-Complex | Early Auditory Cortex | Somatomotor Network | SMT |
| -61 | -15 | -4 | Left-Auditory-5-Complex | Auditory Association Cortex | Somatomotor Network | SMT |
| -22 | -35 | -17 | Left-ParaHippocampal-Area-1 | Medial Temporal Cortex | Default Mode Network | DMN |
| -32 | -38 | -18 | Left-ParaHippocampal-Area-3 | Medial Temporal Cortex | Visual Network | VIS |
| -54 | -8 | -13 | Left-Area-STSd-anterior | Auditory Association Cortex | Default Mode Network | DMN |
| -53 | -33 | -2 | Left-Area-STSd-posterior | Auditory Association Cortex | Default Mode Network | DMN |
| -55 | -36 | -7 | Left-Area-STSv-posterior | Auditory Association Cortex | Default Mode Network | DMN |
| -60 | -11 | -23 | Left-Area-TE1-anterior | Lateral Temporal Cortex | Default Mode Network | DMN |
| -60 | -46 | -13 | Left-Area-TE1-posterior | Lateral Temporal Cortex | Default Mode Network | DMN |
| -58 | -57 | -1 | Left-Area-PHT | Lateral Temporal Cortex | Dorsal Attention Network | DAN |
| -47 | -60 | -11 | Left-Area-PH | MT+ Complex and Neighboring Visual Areas | Dorsal Attention Network | DAN |
| -55 | -45 | 7 | Left-Area-TemporoParietoOccipital-Junction-1 | Temporo-Parieto-Occipital Junction | Somatomotor Network | SMT |

| | | | | | | |
|---|---|---|---|---|---|---|
| -49 | -61 | 10 | Left-Area-TemporoParietoOccipital-Junction-2 | Temporo-Parieto-Occipital Junction | Dorsal Attention Network | DAN |
| -44 | -71 | 15 | Left-Area-TemporoParietoOccipital-Junction-3 | Temporo-Parieto-Occipital Junction | Dorsal Attention Network | DAN |
| -18 | -71 | 28 | Left-Dorsal-Transitional-Visual-Area | Posterior Cingulate Cortex | Visual Network | VIS |
| -40 | -81 | 21 | Left-Area-PGp | Inferior Parietal Cortex | Dorsal Attention Network | DAN |
| -40 | -51 | 38 | Left-Area-IntraParietal-2 | Inferior Parietal Cortex | Frontoparietal Network | FPT |
| -31 | -69 | 39 | Left-Area-IntraParietal-1 | Inferior Parietal Cortex | Frontoparietal Network | FPT |
| -32 | -76 | 24 | Left-Area-IntraParietal-0 | Inferior Parietal Cortex | Dorsal Attention Network | DAN |
| -61 | -24 | 25 | Left-Area-PF-opercular | Inferior Parietal Cortex | Ventral Attention Network | VAN |
| -57 | -39 | 36 | Left-Area-PFm-Complex | Inferior Parietal Cortex | Ventral Attention Network | VAN |
| -48 | -58 | 39 | Left-Area-PFm-Complex | Inferior Parietal Cortex | Default Mode Network | DMN |
| -44 | -59 | 22 | Left-Area-PGi | Inferior Parietal Cortex | Default Mode Network | DMN |
| -40 | -73 | 35 | Left-Area-PGs | Inferior Parietal Cortex | Default Mode Network | DMN |
| -19 | -85 | 37 | Left-Area-V6A | Dorsal Strem Visual Cortex | Visual Network | VIS |
| -19 | -53 | -8 | Left-VentroMedial-Visual-Area-1 | Ventral Stream Visual Cortex | Visual Network | VIS |
| -29 | -60 | -11 | Left-VentroMedial-Visual-Area-3 | Ventral Stream Visual Cortex | Visual Network | VIS |
| -31 | -36 | -15 | Left-ParaHippocampal-Area-2 | Medial Temporal Cortex | Visual Network | VIS |
| -44 | -77 | -2 | Left-Area-V4t | MT+ Complex and Neighboring Visual Areas | Visual Network | VIS |
| -47 | -65 | -2 | Left-Area-FST | MT+ Complex and Neighboring Visual Areas | Dorsal Attention Network | DAN |
| -35 | -86 | 11 | Left-Area-V3CD | MT+ Complex and Neighboring Visual Areas | Visual Network | VIS |
| -46 | -79 | 9 | Left-Area-Lateral-Occipital-3 | MT+ Complex and Neighboring Visual Areas | Visual Network | VIS |
| -28 | -53 | -7 | Left-VentroMedial-Visual-Area-2 | Ventral Stream Visual Cortex | Visual Network | VIS |
| -11 | -52 | 34 | Left-Area-31pd | Posterior Cingulate Cortex | Default Mode Network | DMN |
| -6 | -37 | 42 | Left-Area-31a | Posterior Cingulate Cortex | Default Mode Network | DMN |
| -31 | -52 | -19 | Left-Ventral-Visual-Complex | Ventral Stream Visual Cortex | Visual Network | VIS |
| -5 | 23 | -13 | Left-Area-25 | Anterior Cingulate and Medial Prefrontal Cortex | Limbic Network | LMB |
| -7 | 34 | -14 | Left-Area-s32 | Anterior Cingulate and Medial Prefrontal Cortex | Limbic Network | LMB |
| -38 | -13 | -5 | Left-Area-Posterior-Insular-1 | Insular and Frontal Opercular Cortex | Ventral Attention Network | VAN |
| -36 | -16 | 13 | Left-Insular-Granular-Complex | Insular and Frontal Opercular Cortex | Somatomotor Network | SMT |
| -35 | 26 | 5 | Left-Area-Frontal-Opercular-5 | Insular and Frontal Opercular Cortex | Ventral Attention Network | VAN |

| | | | | | | |
|---|---|---|---|---|---|---|
| -22 | 59 | 4 | Left-Area-posterior-10p | Orbital and Polar Frontal Cortex | Default Mode Network | DMN |
| -43 | 40 | 0 | Left-Area-posterior-47r | Inferior Frontal Cortex | Frontoparietal Network | FPT |
| -45 | -18 | 2 | Left-Medial-Belt-Complex | Early Auditory Cortex | Somatomotor Network | SMT |
| -46 | -27 | 6 | Left-Lateral-Belt-Complex | Early Auditory Cortex | Somatomotor Network | SMT |
| -63 | -22 | 5 | Left-Auditory-4-Complex | Auditory Association Cortex | Somatomotor Network | SMT |
| -52 | -13 | -18 | Left-Area-STSv-anterior | Auditory Association Cortex | Default Mode Network | DMN |
| -44 | -5 | -16 | Left-Para-Insular-Area | Insular and Frontal Opercular Cortex | Ventral Attention Network | VAN |
| -10 | 29 | 27 | Left-Area-anterior-32-prime | Anterior Cingulate and Medial Prefrontal Cortex | Frontoparietal Network | FPT |
| -4 | 35 | 15 | Left-Area-posterior-24 | Anterior Cingulate and Medial Prefrontal Cortex | Default Mode Network | DMN |
| 12 | -81 | 2 | Right-Primary-Visual-Cortex | Primary Visual Cortex | Visual Network | VIS |
| 43 | -66 | 3 | Right-Medial-Superior-Temporal-Area | MT+ Complex and Neighboring Visual Areas | Visual Network | VIS |
| 17 | -77 | 29 | Right-Sixth-Visual-Area | Dorsal Strem Visual Cortex | Visual Network | VIS |
| 13 | -79 | 3 | Right-Second-Visual-Area | Early Visual Cortex | Visual Network | VIS |
| 19 | -85 | 6 | Right-Third-Visual-Area | Early Visual Cortex | Visual Network | VIS |
| 31 | -84 | -3 | Right-Fourth-Visual-Area | Early Visual Cortex | Visual Network | VIS |
| 31 | -71 | -13 | Right-Eighth-Visual-Area | Ventral Stream Visual Cortex | Visual Network | VIS |
| 29 | -17 | 54 | Right-Primary-Motor-Cortex | Somatosensory and Motor Cortex | Somatomotor Network | SMT |
| 37 | -22 | 51 | Right-Primary-Sensory-Cortex | Somatosensory and Motor Cortex | Somatomotor Network | SMT |
| 42 | -3 | 49 | Right-Frontal-Eye-Fields | Premotor Cortex | Dorsal Attention Network | DAN |
| 45 | 2 | 36 | Right-Premotor-Eye-Field | Premotor Cortex | Dorsal Attention Network | DAN |
| 49 | 0 | 45 | Right-Area-55b | Premotor Cortex | Ventral Attention Network | VAN |
| 16 | -87 | 29 | Right-Area-V3A | Dorsal Strem Visual Cortex | Visual Network | VIS |
| 6 | -37 | 20 | Right-RetroSplenial-Complex | Posterior Cingulate Cortex | Default Mode Network | DMN |
| 11 | -69 | 36 | Right-Parieto-Occipital-Sulcus-Area-2 | Posterior Cingulate Cortex | Frontoparietal Network | FPT |
| 25 | -81 | 30 | Right-Seventh-Visual-Area | Dorsal Strem Visual Cortex | Visual Network | VIS |
| 25 | -69 | 35 | Right-IntraParietal-Sulcus-Area-1 | Dorsal Strem Visual Cortex | Dorsal Attention Network | DAN |
| 39 | -55 | -19 | Right-Fusiform-Face-Complex | Ventral Stream Visual Cortex | Visual Network | VIS |
| 29 | -75 | 20 | Right-Area-V3B | Dorsal Strem Visual Cortex | Visual Network | VIS |
| 39 | -83 | 3 | Right-Area-Lateral-Occipital-1 | MT+ Complex and Neighboring Visual Areas | Visual Network | VIS |

| | | | | | | |
|---|---|---|---|---|---|---|
| 44 | -82 | -5 | Right-Area-Lateral-Occipital-2 | MT+ Complex and Neighboring Visual Areas | Visual Network | VIS |
| 43 | -79 | -14 | Right-Posterior-InferoTemporal-Complex | Ventral Stream Visual Cortex | Visual Network | VIS |
| 47 | -72 | 6 | Right-Middle-Temporal-Area | MT+ Complex and Neighboring Visual Areas | Visual Network | VIS |
| 44 | -23 | 10 | Right-Primary-Auditory-Cortex | Early Auditory Cortex | Somatomotor Network | SMT |
| 61 | -39 | 21 | Right-PeriSylvian-Language-Area | Temporo-Parieto-Occipital Junction | Ventral Attention Network | VAN |
| 9 | 16 | 63 | Right-Superior-Frontal-Language-Area | DorsoLateral Prefrontal Cortex | Default Mode Network | DMN |
| 7 | -52 | 49 | Right-PreCuneus-Visual-Area | Posterior Cingulate Cortex | Dorsal Attention Network | DAN |
| 57 | -42 | 14 | Right-Superior-Temporal-Visual-Area | Temporo-Parieto-Occipital Junction | Ventral Attention Network | VAN |
| 6 | -66 | 49 | Right-Medial-Area-7P | Superior Parietal Cortex | Frontoparietal Network | FPT |
| 5 | -60 | 33 | Right-Area-7m | Posterior Cingulate Cortex | Default Mode Network | DMN |
| 11 | -58 | 15 | Right-Parieto-Occipital-Sulcus-Area-1 | Posterior Cingulate Cortex | Default Mode Network | DMN |
| 4 | -23 | 38 | Right-Area-23d | Posterior Cingulate Cortex | Default Mode Network | DMN |
| 5 | -53 | 18 | Right-Area-ventral-23-a+b | Posterior Cingulate Cortex | Default Mode Network | DMN |
| 4 | -41 | 31 | Right-Area-dorsal-23-a+b | Posterior Cingulate Cortex | Default Mode Network | DMN |
| 8 | -45 | 33 | Right-Area-31p-ventral | Posterior Cingulate Cortex | Default Mode Network | DMN |
| 5 | -38 | 63 | Right-Area-5m | Paracentral Lobular and Mid Cingulate Cortex | Somatomotor Network | SMT |
| 13 | -38 | 51 | Right-Area-5m-ventral | Paracentral Lobular and Mid Cingulate Cortex | Ventral Attention Network | VAN |
| 11 | -32 | 42 | Right-Area-23c | Paracentral Lobular and Mid Cingulate Cortex | Ventral Attention Network | VAN |
| 12 | -44 | 71 | Right-Area-5L | Paracentral Lobular and Mid Cingulate Cortex | Somatomotor Network | SMT |
| 8 | -18 | 50 | Right-Dorsal-Area-24d | Paracentral Lobular and Mid Cingulate Cortex | Somatomotor Network | SMT |
| 11 | -5 | 46 | Right-Ventral-Area-24d | Paracentral Lobular and Mid Cingulate Cortex | Somatomotor Network | SMT |
| 20 | -52 | 64 | Right-Lateral-Area-7A | Superior Parietal Cortex | Dorsal Attention Network | DAN |
| 7 | 4 | 57 | Right-Supplementary-and-Cingulate-Eye-Field | Paracentral Lobular and Mid Cingulate Cortex | Ventral Attention Network | VAN |
| 18 | 4 | 64 | Right-Area-6m-anterior | Paracentral Lobular and Mid Cingulate Cortex | Ventral Attention Network | VAN |
| 9 | -60 | 60 | Right-Medial-Area-7A | Superior Parietal Cortex | Dorsal Attention Network | DAN |
| 13 | -70 | 54 | Right-Lateral-Area-7P | Superior Parietal Cortex | Dorsal Attention Network | DAN |
| 33 | -49 | 61 | Right-Area-7PC | Superior Parietal Cortex | Dorsal Attention Network | DAN |
| 28 | -54 | 53 | Right-Area-Lateral-IntraParietal-ventral | Superior Parietal Cortex | Dorsal Attention Network | DAN |
| 19 | -61 | 61 | Right-Ventral-IntraParietal-Complex | Superior Parietal Cortex | Dorsal Attention Network | DAN |

| | | | | | | |
|---|---|---|---|---|---|---|
| 24 | -64 | 45 | Right-Medial-IntraParietal-Area | Superior Parietal Cortex | Dorsal Attention Network | DAN |
| 45 | -24 | 54 | Right-Area-1 | Somatosensory and Motor Cortex | Somatomotor Network | SMT |
| 37 | -32 | 51 | Right-Area-2 | Somatosensory and Motor Cortex | Somatomotor Network | SMT |
| 31 | -21 | 45 | Right-Area-3a | Somatosensory and Motor Cortex | Somatomotor Network | SMT |
| 36 | -12 | 62 | Right-Dorsal-area-6 | Premotor Cortex | Somatomotor Network | SMT |
| 15 | -12 | 66 | Right-Area-6mp | Paracentral Lobular and Mid Cingulate Cortex | Somatomotor Network | SMT |
| 58 | 4 | 29 | Right-Ventral-Area-6 | Premotor Cortex | Somatomotor Network | SMT |
| 5 | -2 | 40 | Right-Area-Posterior-24-prime | Anterior Cingulate and Medial Prefrontal Cortex | Ventral Attention Network | VAN |
| 4 | 11 | 30 | Right-Area-33-prime | Anterior Cingulate and Medial Prefrontal Cortex | Ventral Attention Network | VAN |
| 5 | 18 | 32 | Right-Anterior-24-prime | Anterior Cingulate and Medial Prefrontal Cortex | Ventral Attention Network | VAN |
| 11 | 14 | 39 | Right-Area-p32-prime | Anterior Cingulate and Medial Prefrontal Cortex | Ventral Attention Network | VAN |
| 6 | 38 | 0 | Right-Area-a24 | Anterior Cingulate and Medial Prefrontal Cortex | Default Mode Network | DMN |
| 10 | 37 | 24 | Right-Area-dorsal-32 | Anterior Cingulate and Medial Prefrontal Cortex | Default Mode Network | DMN |
| 6 | 28 | 46 | Right-Area-8BM | Anterior Cingulate and Medial Prefrontal Cortex | Frontoparietal Network | FPT |
| 11 | 44 | -2 | Right-Area-p32 | Anterior Cingulate and Medial Prefrontal Cortex | Default Mode Network | DMN |
| 8 | 47 | -8 | Right-Area-10r | Anterior Cingulate and Medial Prefrontal Cortex | Default Mode Network | DMN |
| 34 | 30 | -16 | Right-Area-47m | Orbital and Polar Frontal Cortex | Default Mode Network | DMN |
| 38 | 20 | 47 | Right-Area-8Av | DorsoLateral Prefrontal Cortex | Frontoparietal Network | FPT |
| 23 | 27 | 44 | Right-Area-8Ad | DorsoLateral Prefrontal Cortex | Default Mode Network | DMN |
| 8 | 52 | 22 | Right-Area-9-Middle | Anterior Cingulate and Medial Prefrontal Cortex | Default Mode Network | DMN |
| 11 | 38 | 49 | Right-Area-8B-Lateral | DorsoLateral Prefrontal Cortex | Default Mode Network | DMN |
| 18 | 47 | 34 | Right-Area-9-Posterior | DorsoLateral Prefrontal Cortex | Default Mode Network | DMN |
| 11 | 64 | 6 | Right-Area-10d | Orbital and Polar Frontal Cortex | Default Mode Network | DMN |
| 39 | 18 | 35 | Right-Area-8C | DorsoLateral Prefrontal Cortex | Frontoparietal Network | FPT |
| 52 | 17 | 12 | Right-Area-44 | Inferior Frontal Cortex | Frontoparietal Network | FPT |
| 50 | 27 | 3 | Right-Area-45 | Inferior Frontal Cortex | Default Mode Network | DMN |
| 39 | 48 | -7 | Right-Area-anterior-47r | Inferior Frontal Cortex | Frontoparietal Network | FPT |
| 50 | 8 | 16 | Right-Rostral-Area-6 | Premotor Cortex | Ventral Attention Network | VAN |
| 41 | 18 | 22 | Right-Area-IFJa | Inferior Frontal Cortex | Frontoparietal Network | FPT |

| | | | | | | |
|---|---|---|---|---|---|---|
| 37 | 8 | 27 | Right-Area-IFJp | Inferior Frontal Cortex | Dorsal Attention Network | DAN |
| 47 | 27 | 17 | Right-Area-IFSp | Inferior Frontal Cortex | Frontoparietal Network | FPT |
| 47 | 34 | 6 | Right-Area-IFSa | Inferior Frontal Cortex | Frontoparietal Network | FPT |
| 44 | 31 | 28 | Right-Area-posterior-9-46v | DorsoLateral Prefrontal Cortex | Frontoparietal Network | FPT |
| 36 | 37 | 27 | Right-Area-46 | DorsoLateral Prefrontal Cortex | Frontoparietal Network | FPT |
| 39 | 49 | 11 | Right-Area-anterior-9-46v | DorsoLateral Prefrontal Cortex | Frontoparietal Network | FPT |
| 29 | 45 | 23 | Right-Area-9-46d | DorsoLateral Prefrontal Cortex | Frontoparietal Network | FPT |
| 18 | 58 | 21 | Right-Area-9-anterior | DorsoLateral Prefrontal Cortex | Default Mode Network | DMN |
| 5 | 49 | -15 | Right-Area-10v | Anterior Cingulate and Medial Prefrontal Cortex | Limbic Network | LMB |
| 25 | 58 | -7 | Right-Area-anterior-10p | Orbital and Polar Frontal Cortex | Frontoparietal Network | FPT |
| 13 | 58 | -16 | Right-Polar-10p | Orbital and Polar Frontal Cortex | Limbic Network | LMB |
| 21 | 24 | -20 | Right-Area-13l | Orbital and Polar Frontal Cortex | Limbic Network | LMB |
| 32 | 21 | -18 | Right-Area-47s | Orbital and Polar Frontal Cortex | Default Mode Network | DMN |
| 30 | -54 | 43 | Right-Area-Lateral-IntraParietal-dorsal | Superior Parietal Cortex | Dorsal Attention Network | DAN |
| 27 | -2 | 51 | Right-Area-6-anterior | Premotor Cortex | Dorsal Attention Network | DAN |
| 35 | 11 | 54 | Right-Inferior-6-8-Transitional-Area | DorsoLateral Prefrontal Cortex | Frontoparietal Network | FPT |
| 20 | 19 | 56 | Right-Superior-6-8-Transitional-Area | DorsoLateral Prefrontal Cortex | Frontoparietal Network | FPT |
| 56 | -2 | 10 | Right-Area-43 | Posterior Opercular Cortex | Somatomotor Network | SMT |
| 58 | -14 | 16 | Right-Area-OP4/PV | Posterior Opercular Cortex | Somatomotor Network | SMT |
| 44 | -22 | 18 | Right-Area-OP1/SII | Posterior Opercular Cortex | Somatomotor Network | SMT |
| 40 | -15 | 18 | Right-Area-OP2-3/VS | Posterior Opercular Cortex | Somatomotor Network | SMT |
| 39 | -23 | 2 | Right-Area-52 | Early Auditory Cortex | Somatomotor Network | SMT |
| 42 | -33 | 17 | Right-RetroInsular-Cortex | Early Auditory Cortex | Somatomotor Network | SMT |
| 48 | -30 | 22 | Right-Area-PFcm | Early Auditory Cortex | Somatomotor Network | SMT |
| 39 | -5 | -1 | Right-Posterior-Insular-Area-2 | Insular and Frontal Opercular Cortex | Ventral Attention Network | VAN |
| 52 | -1 | -7 | Right-Area-TA2 | Auditory Association Cortex | Somatomotor Network | SMT |
| 40 | 13 | 7 | Right-Frontal-Opercular-Area-4 | Insular and Frontal Opercular Cortex | Ventral Attention Network | VAN |
| 38 | 8 | 1 | Right-Middle-Insular-Area | Insular and Frontal Opercular Cortex | Ventral Attention Network | VAN |
| 34 | 5 | -18 | Right-Pirform-Cortex | Insular and Frontal Opercular Cortex | Ventral Attention Network | VAN |

| | | | | | | |
|---|---|---|---|---|---|---|
| 34 | 23 | -3 | Right-Anterior-Ventral-Insular-Area | Insular and Frontal Opercular Cortex | Frontoparietal Network | FPT |
| 35 | 13 | -12 | Right-Anterior-Agranular-Insula-Complex | Insular and Frontal Opercular Cortex | Ventral Attention Network | VAN |
| 48 | 3 | 5 | Right-Frontal-Opercular-Area-1 | Posterior Opercular Cortex | Ventral Attention Network | VAN |
| 35 | 6 | 11 | Right-Frontal-Opercular-Area-3 | Insular and Frontal Opercular Cortex | Ventral Attention Network | VAN |
| 41 | -4 | 15 | Right-Frontal-Opercular-Area-2 | Insular and Frontal Opercular Cortex | Somatomotor Network | SMT |
| 52 | -27 | 40 | Right-Area-PFt | Inferior Parietal Cortex | Dorsal Attention Network | DAN |
| 37 | -41 | 42 | Right-Anterior-IntraParietal-Area | Superior Parietal Cortex | Dorsal Attention Network | DAN |
| 18 | -35 | -8 | Right-PreSubiculum | Medial Temporal Cortex | Visual Network | VIS |
| 20 | -51 | 1 | Right-ProStriate-Area | Posterior Cingulate Cortex | Visual Network | VIS |
| 51 | 10 | -19 | Right-Area-STGa | Auditory Association Cortex | Default Mode Network | DMN |
| 57 | -18 | 5 | Right-ParaBelt-Complex | Early Auditory Cortex | Somatomotor Network | SMT |
| 61 | -17 | -3 | Right-Auditory-5-Complex | Auditory Association Cortex | Somatomotor Network | SMT |
| 23 | -34 | -17 | Right-ParaHippocampal-Area-1 | Medial Temporal Cortex | Visual Network | VIS |
| 53 | -7 | -14 | Right-Area-STSd-anterior | Auditory Association Cortex | Default Mode Network | DMN |
| 49 | -29 | -3 | Right-Area-STSd-posterior | Auditory Association Cortex | Default Mode Network | DMN |
| 59 | -33 | -7 | Right-Area-STSv-posterior | Auditory Association Cortex | Default Mode Network | DMN |
| 60 | -7 | -25 | Right-Area-TE1-anterior | Lateral Temporal Cortex | Default Mode Network | DMN |
| 61 | -44 | -13 | Right-Area-TE1-posterior | Lateral Temporal Cortex | Frontoparietal Network | FPT |
| 59 | -53 | -4 | Right-Area-PHT | Lateral Temporal Cortex | Dorsal Attention Network | DAN |
| 49 | -62 | -11 | Right-Area-PH | MT+ Complex and Neighboring Visual Areas | Dorsal Attention Network | DAN |
| 54 | -43 | 7 | Right-Area-TemporoParietoOccipital-Junction-1 | Temporo-Parieto-Occipital Junction | Ventral Attention Network | VAN |
| 54 | -58 | 7 | Right-Area-TemporoParietoOccipital-Junction-2 | Temporo-Parieto-Occipital Junction | Dorsal Attention Network | DAN |
| 41 | -64 | 14 | Right-Area-TemporoParietoOccipital-Junction-3 | Temporo-Parieto-Occipital Junction | Dorsal Attention Network | DAN |
| 20 | -67 | 28 | Right-Dorsal-Transitional-Visual-Area | Posterior Cingulate Cortex | Visual Network | VIS |
| 41 | -79 | 23 | Right-Area-PGp | Inferior Parietal Cortex | Dorsal Attention Network | DAN |
| 43 | -45 | 41 | Right-Area-IntraParietal-2 | Inferior Parietal Cortex | Frontoparietal Network | FPT |

| | | | | | | |
|---|---|---|---|---|---|---|
| 35 | -66 | 42 | Right-Area-IntraParietal-1 | Inferior Parietal Cortex | Frontoparietal Network | FPT |
| 33 | -73 | 27 | Right-Area-IntraParietal-0 | Inferior Parietal Cortex | Dorsal Attention Network | DAN |
| 60 | -21 | 27 | Right-Area-PF-opercular | Inferior Parietal Cortex | Ventral Attention Network | VAN |
| 58 | -34 | 36 | Right-Area-PFm-Complex | Inferior Parietal Cortex | Ventral Attention Network | VAN |
| 52 | -50 | 39 | Right-Area-PFm-Complex | Inferior Parietal Cortex | Frontoparietal Network | FPT |
| 47 | -59 | 23 | Right-Area-PGi | Inferior Parietal Cortex | Default Mode Network | DMN |
| 43 | -67 | 38 | Right-Area-PGs | Inferior Parietal Cortex | Default Mode Network | DMN |
| 22 | -81 | 41 | Right-Area-V6A | Dorsal Strem Visual Cortex | Visual Network | VIS |
| 19 | -53 | -8 | Right-VentroMedial-Visual-Area-1 | Ventral Stream Visual Cortex | Visual Network | VIS |
| 29 | -59 | -10 | Right-VentroMedial-Visual-Area-3 | Ventral Stream Visual Cortex | Visual Network | VIS |
| 32 | -35 | -15 | Right-ParaHippocampal-Area-2 | Medial Temporal Cortex | Visual Network | VIS |
| 45 | -78 | -2 | Right-Area-V4t | MT+ Complex and Neighboring Visual Areas | Visual Network | VIS |
| 46 | -64 | -3 | Right-Area-FST | MT+ Complex and Neighboring Visual Areas | Dorsal Attention Network | DAN |
| 34 | -83 | 14 | Right-Area-V3CD | MT+ Complex and Neighboring Visual Areas | Visual Network | VIS |
| 46 | -77 | 9 | Right-Area-Lateral-Occipital-3 | MT+ Complex and Neighboring Visual Areas | Visual Network | VIS |
| 29 | -54 | -7 | Right-VentroMedial-Visual-Area-2 | Ventral Stream Visual Cortex | Visual Network | VIS |
| 12 | -52 | 35 | Right-Area-31pd | Posterior Cingulate Cortex | Default Mode Network | DMN |
| 6 | -42 | 41 | Right-Area-31a | Posterior Cingulate Cortex | Frontoparietal Network | FPT |
| 30 | -48 | -19 | Right-Ventral-Visual-Complex | Ventral Stream Visual Cortex | Visual Network | VIS |
| 5 | 20 | -14 | Right-Area-25 | Anterior Cingulate and Medial Prefrontal Cortex | Limbic Network | LMB |
| 6 | 33 | -13 | Right-Area-s32 | Anterior Cingulate and Medial Prefrontal Cortex | Default Mode Network | DMN |
| 39 | -13 | -4 | Right-Area-Posterior-Insular-1 | Insular and Frontal Opercular Cortex | Ventral Attention Network | VAN |
| 36 | -15 | 14 | Right-Insular-Granular-Complex | Insular and Frontal Opercular Cortex | Somatomotor Network | SMT |
| 37 | 26 | 5 | Right-Area-Frontal-Opercular-5 | Insular and Frontal Opercular Cortex | Ventral Attention Network | VAN |
| 25 | 58 | 5 | Right-Area-posterior-10p | Orbital and Polar Frontal Cortex | Frontoparietal Network | FPT |
| 45 | 41 | -3 | Right-Area-posterior-47r | Inferior Frontal Cortex | Frontoparietal Network | FPT |
| 45 | -19 | 4 | Right-Medial-Belt-Complex | Early Auditory Cortex | Somatomotor Network | SMT |
| 49 | -26 | 8 | Right-Lateral-Belt-Complex | Early Auditory Cortex | Somatomotor Network | SMT |
| 64 | -17 | 4 | Right-Auditory-4-Complex | Auditory Association Cortex | Somatomotor Network | SMT |

| | | | | | | |
|---|---|---|---|---|---|---|
| 56 | -14 | -17 | Right-Area-STSv-anterior | Auditory Association Cortex | Default Mode Network | DMN |
| 63 | -27 | -16 | Right-Area-TE1-Middle | Lateral Temporal Cortex | Default Mode Network | DMN |
| 45 | -7 | -13 | Right-Para-Insular-Area | Insular and Frontal Opercular Cortex | Ventral Attention Network | VAN |
| 10 | 28 | 28 | Right-Area-anterior-32-prime | Anterior Cingulate and Medial Prefrontal Cortex | Frontoparietal Network | FPT |
| 5 | 35 | 16 | Right-Area-posterior-24 | Anterior Cingulate and Medial Prefrontal Cortex | Default Mode Network | DMN |

### Figures

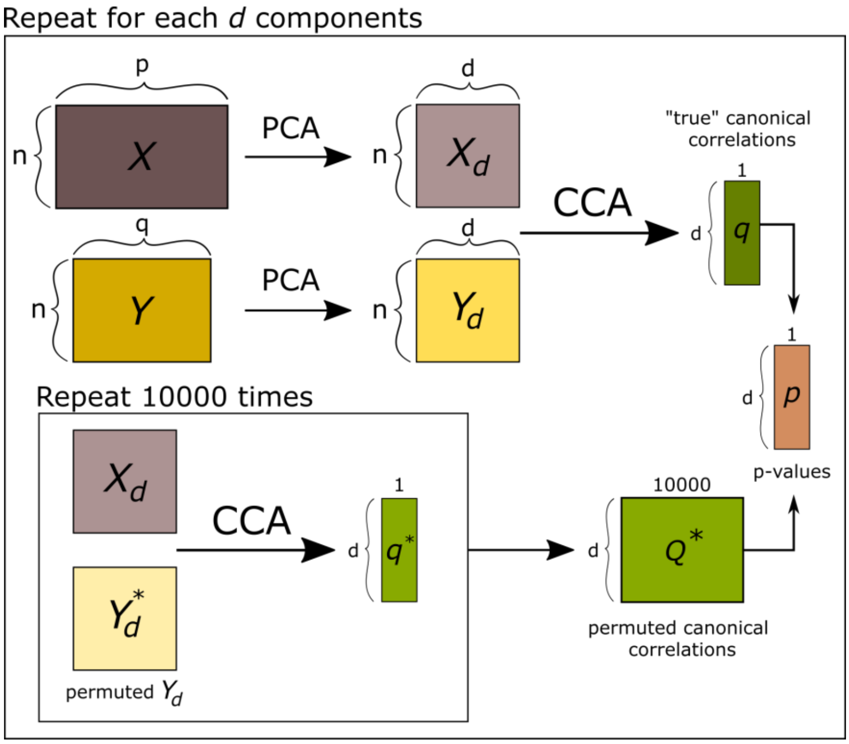


**Figure S1:** Permutation framework used to jointly find the optimal number of PCA components and estimate the statistically significant CCA modes.

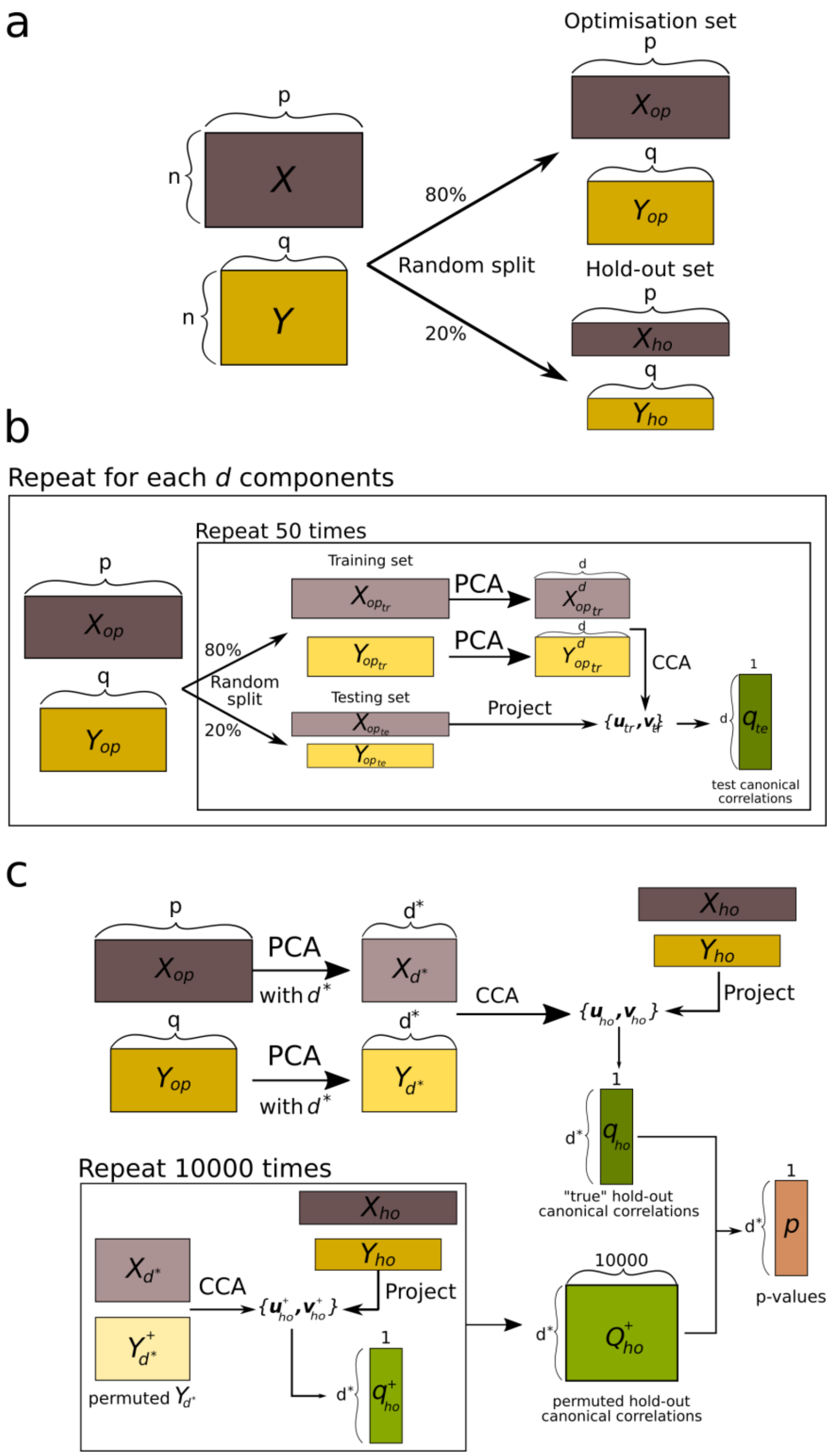


**Figure S2:** Hold-out framework used to jointly find the optimal number of PCA components and estimate the statistically significant CCA modes. **(a)** Random split of the data into optimisation and hold-out sets; **(b)** optimisation of the number of PCA components; **(c)** validation of the model using a hold-out set.

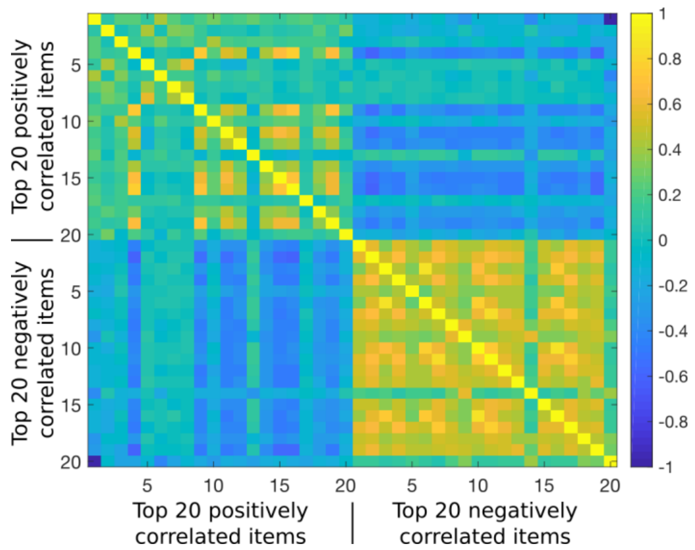


**Figure S3:** Correlations between the top 20 positive and top 20 negative behavioural items of the first CCA mode. Male gender (first item) is weakly associated with the other positive behavioural items (items 2-19 with mean correlation=0.20). Female gender (last item) is weakly associated with the other negative behavioural items (items 1-19 with mean correlation=0.17).

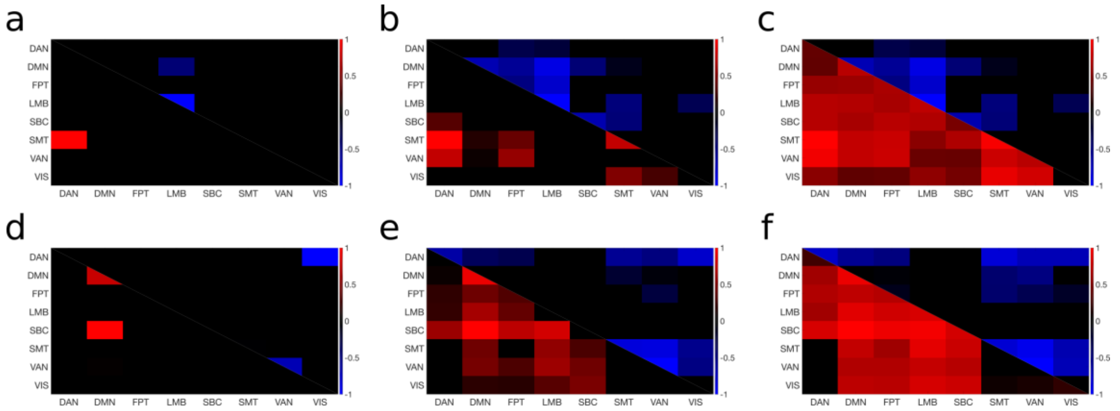


**Figure S4:** Mean correlations between and within resting-state networks for the first (**a-c**) and the second (**d-f**) CCA mode at three different levels of top connections: top 20 (**a, d**), top 0.5% (**b, e**) and top 5% (**c, f**) of most positively/negatively correlated connections. Positive correlations (red) and negative correlations (blue) are summarized separately in the lower and upper triangular matrices, respectively. The mean absolute correlations are log-transformed and normalized for easier comparison between the three levels. Dorsal Attention Network (DAN); Default Mode Network (DMN); Frontoparietal Network (FPT); Limbic Network (LMB); Subcortex (SBC); Somatomotor Network (SMT); Ventral Attention Network (VAN); Visual Network (VIS).

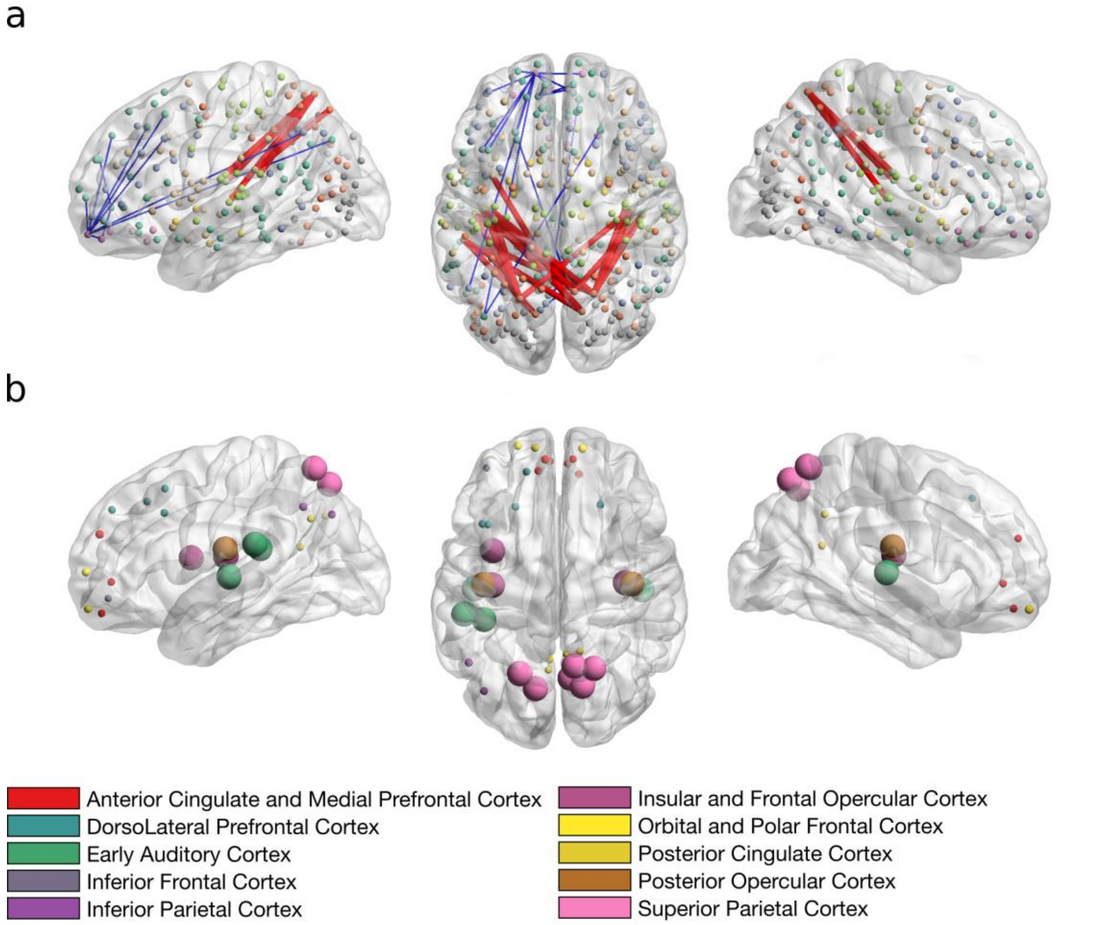


**Figure S5:** Correlations between the brain connectivity variables and the brain canonical variate (brain scores of all subjects) of the first CCA mode in sagittal (left and right) and axial views (middle). Notations are as in Fig. 4 of the main text except that nodes are colour coded by gross anatomical regions used in Glasser et al 2016[17]. The full list of correlation values and respective labels can be found in Supplementary Table S3.

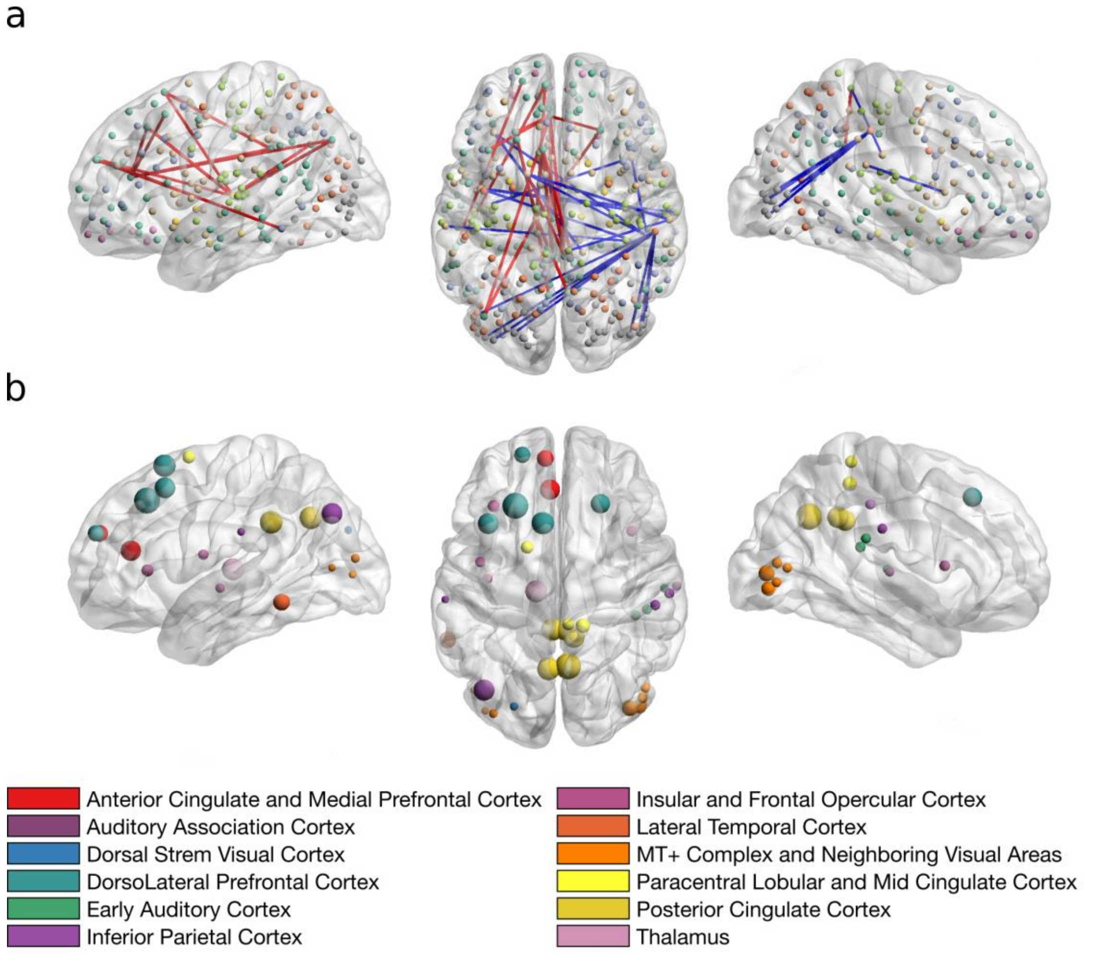


**Figure S6:** Correlations between the brain connectivity variables and the brain canonical variate (brain scores of all subjects) of the second CCA mode in sagittal (left and right) and axial views (middle). Notations are as in Fig. 5 of the main text except that nodes are colour coded by gross anatomical regions used in Glasser et al 2016[17]. The full list of correlation values and respective labels can be found in Supplementary Table S4.

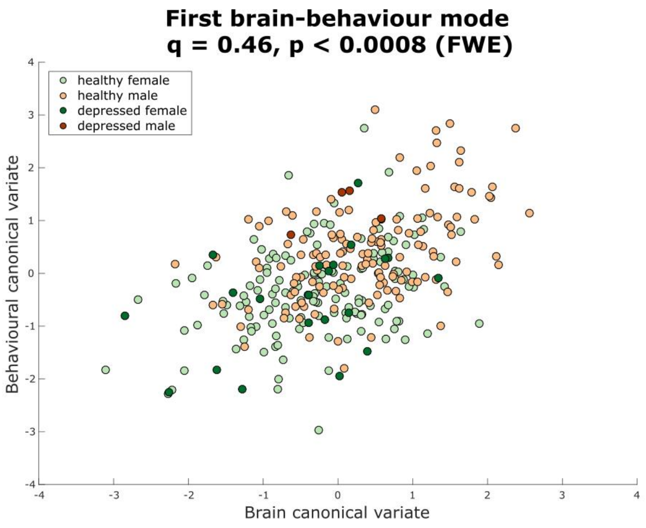


**Figure S7:** Significant brain-behaviour mode of covariation using the multiple hold-out framework. Scatter plot showing the brain and behaviour scores for the first CCA mode, where each dot represents an individual subject. Subjects are colour coded by gender and clinical diagnosis. The canonical hold-out correlation, *q*, and corresponding p-value are shown on the top of the plot.

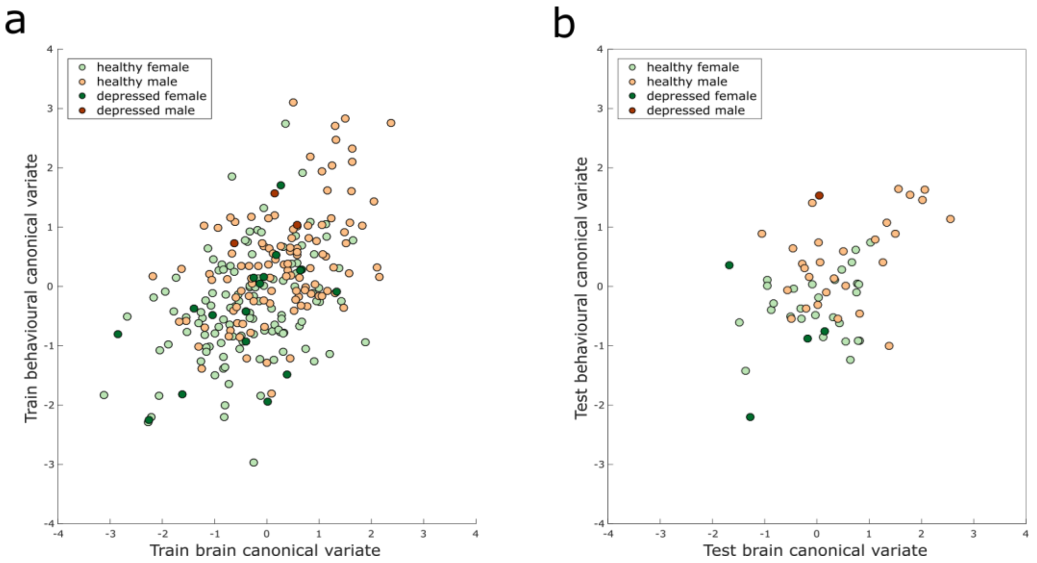


**Figure S8:** Significant brain-behaviour mode of population covariation for the training (**a**) and testing set (**b**) using the multiple hold-out framework. All the conventions are as in Supplementary Fig. S7.

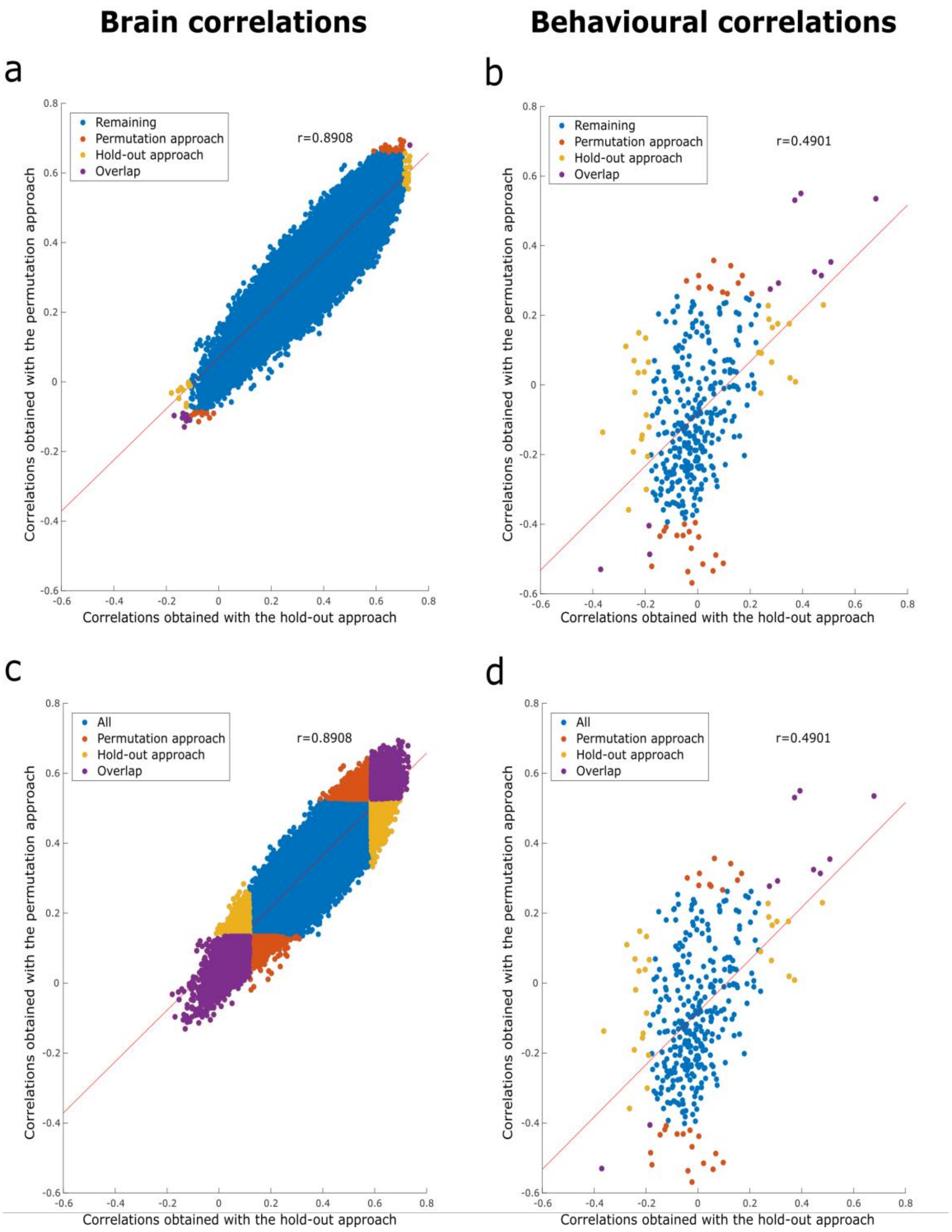


**Figure S9:** Scatter plots showing the brain (**a,c**) and behaviour (**b,d**) correlations with the first CCA mode obtained with both frameworks. (**a,b**) The overlap (purple) between the top 20 most positively/negatively correlated variables obtained with the permutation (orange) and multiple hold-out framework (yellow) is shown; (**c,d**) the same colour scheme is used to show the overlap between the top 5% most positively/negatively correlated variables obtained with the permutation and multiple hold-out framework. Blue denotes the remaining variables.
